# Proteostasis and Energetics as Proteome Hallmarks of Aging and Influenza Challenge in Pulmonary Disease

**DOI:** 10.1101/769737

**Authors:** Salvatore Loguercio, Darren M. Hutt, Alexandre Rosa Campos, Thomas Stoeger, Rogan A. Grant, Alexandra C McQuattie-Pimentel, Hiam Abdala-Valencia, Ziyan Lu, Nikita Joshi, Karen Ridge, Navdeep S Chandel, Jacob I. Sznajder, Richard I Morimoto, Alexander V. Misharin, G.R. Scott Budinger, William E. Balch

## Abstract

Aging is associated with an increased risk for the development of many diseases. This is exemplified by the increased incidence of lung injury, muscle dysfunction and cognitive impairment in the elderly following influenza infection. Because the infectious cycle of flu is dependent upon the properties of the host, we examined the proteome of <u>alveolar</u> macrophages (AM) and type 2 cells (AT2) obtained from young (3 months) and old (18 months) naïve mice and mice exposed to influenza A. Our proteomics data show that there is a maladaptive collapse of the proteostasis network (PN) and changes in mitochondrial pathways in the aged naïve AM and AT2 proteomes. The mitochondrial imbalance and proteostatic collapse seen in aged cells places an excessive folding burden on these cells, which is further exacerbated following exposure to influenza A. Specifically, we see an imbalance in Hsp70 co-chaperones involved in protein folding and Hsp90 co-chaperones important for stress signaling pathways that are essential for cellular protection during aging. The acute challenge of influenza A infection of young and aged AM and AT2 cells reveals that age-associated changes in the chaperome affect the ability of these cells to properly manage the infection and post-infection biology, contributing to cytotoxicity. We posit that proteomic profiling of individual cell type specific responses provides a high impact approach to pinpoint fundamental molecular relationships that may contribute to the susceptibility to aging and environmental stress, providing a platform to identify new targets for therapeutic intervention to improve resiliency in the elderly.

## Introduction

As one of the first lines of defense of the organism, the lung is exposed to many environmental stressors that can challenge its functionality (1–6). As such, lung disease is the third leading cause of death in the population. Furthermore, aging related deterioration of pulmonary tissue presents additional challenges including chronic obstructive pulmonary disease (COPD), emphysema and pneumonia. While the pulmonary tissue is capable of surviving influenza-associated challenges, elderly patients have an increased risk of developing age-related disorders including persistent lung injury, skeletal muscle dysfunction leading to immobility, dementia, and cognitive impairment. Although recent transcriptomic and proteomics analyses have begun to highlight differences among the more than 40 cell types that comprise the lungs (3, 7) as well as their respective response to aging (7), the complexity of this organ complicates our ability to ascertain the underlying cause for this increased susceptibility to pneumonia and other diseases (6,8,9).

The long-term health of pulmonary tissue (2,6,8,10–13) is inextricably linked to the sustainability of the protein fold and its function (14–18). Protein homeostasis or proteostasis (18–28), is a collection of integrated biological pathways, referred to as the proteostasis network (PN), that balance protein biosynthesis, folding, trafficking, post-translational modifications and clearance through multiple degradation pathways in response to endogenous and exogenous folding stress during aging (6,16,18–21,23,25–27,29,30). Specifically, proteostasis components involved in protein folding management are referred to as the chaperome (31). These pathways are tightly coupled to the functionality of mitochondria, which manage the energetic health of the cell (32). The health of the mitochondria has received increased attention given its role in managing oxidative stress and intermediary metabolism in the lung (33–37). Recently, the life-span extending drug, metformin, was shown to protect the lung from oxidative stress by inhibiting the activity of the mitochondrial complex I leading to a reduction in reactive oxygen species (ROS) emergent from the complex I machinery (35,38,39). It is now apparent that changes in the cellular transcriptome (3,7,40), proteome (7, 40), stress-related PN pathways (2,6,17,41) and mitochondrial pathways (12,34–36,39,42–46) can have complex effects on the functionality of the lung (19,47–51). However, the nature of the age-associated changes that contribute to the differential sensitivity of young and old lungs to influenza and resulting pneumonia pathophysiology remain enigmatic.

Herein, we have developed a rigorous quantitative mass spectrometry (MS) based workflow to identify age-associated proteomic changes that contribute to the cell-type specific response of alveolar macrophages (AM) and type II (AT2) cells to influenza A infection and the subsequent increased sensitivity of aged lung tissue to developing other age-associated disorders. The MS results reveal significant cell-type specific changes in both mitochondrial and PN pathways impacting the energetics and resiliency of protein folding in the cell. These data lead us to suggest a role for the <u>c</u>o-chaperone <u>a</u>daptive <u>re</u>sponse (CARE) network in managing chronic misfolding in the aging lung and in response to acute challenges to folding by viral infection. The subset of CARE PN components encompasses members of the <u>u</u>nfolded <u>p</u>rotein response (UPR) pathways, the integrated stress response (ISR) linked <u>mito</u>chondrial UPR (UPR_mito_) (ISR-UPR_mito_) pathways, and the Hsf1-mediated heat shock response (HSR). The HSR pathway is charged with managing client delivery to the cytosolic core <u>h</u>eat <u>s</u>hock <u>c</u>ognate (Hsc) 70, <u>h</u>eat <u>s</u>hock <u>p</u>rotein (Hsp) 70 (Hsc/p70) and Hsp90 chaperones that control protein folding and stress responses to misfolding. Understanding the role of proteostasis pathways in lung biology across the population will guide future machine learning efforts to advance our understanding of the spatial relationships (19) that describe the contribution of genetic diversity in the population to resiliency in the individual and aging related stress (52, 53).

## Results

### Proteomic profiling of AT2 and AM lung cells in response to aging and influenza A

Aging is associated with alterations in dynamic biological and physiological processes contributing to human healthspan (54). While some of these biological changes may be benign, others are associated with significant increased risk of disease liability. To distinguish this, we examine the proteome of the aging lung challenged by exposure to influenza A (abbreviated as influenza (F)). We examined isolated <u>a</u>lveolar <u>m</u>acrophages (AM) and <u>a</u>lveolar type <u>II</u> (AT2) cells from the lungs of <u>y</u>oung (Y) (3-4 months) and <u>o</u>ld (O) (18 months) <u>n</u>aïve (N) mice or mice exposed to a survivable dose (10 pfu) of influenza as previously described (35, 55). The proteins from the respective cell lysates were trypsin digested and subjected to 1-<u>d</u>imensional liquid <u>c</u>hromatography (1D-LC) followed by tandem <u>m</u>ass <u>s</u>pectrometry (MS/MS) to characterize the <u>w</u>hole <u>c</u>ell <u>p</u>roteomic (WCP) profile for each condition (**Fig 1**). Our analysis identified 4200 total proteins from AM cells and 4046 total proteins from AT2 cells. A thorough quality control analysis of the WCP datasets was performed to ensure suitability of the data (see Methods) (**Fig S1**).

**Figure 1.**
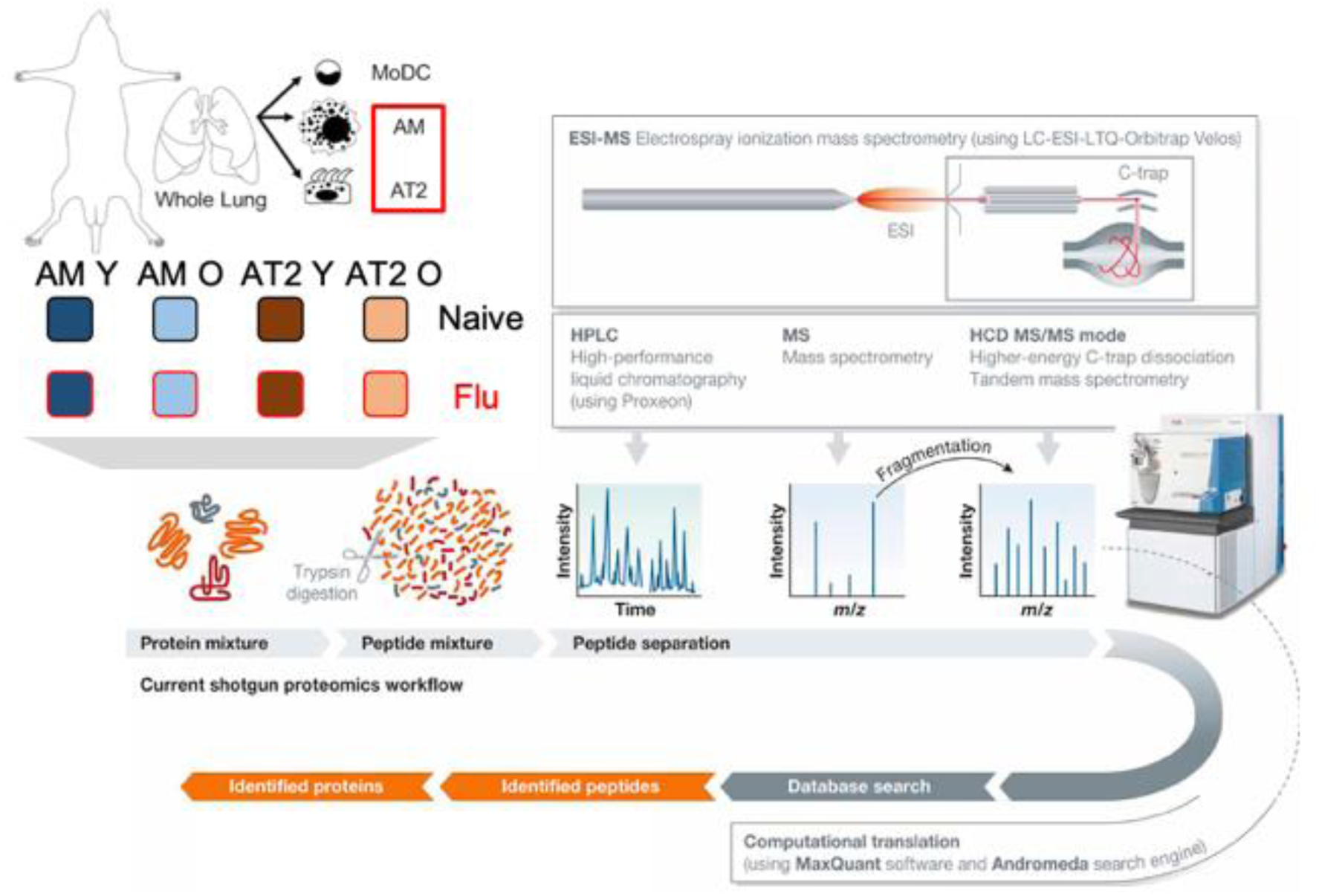
Schematic diagram of the experimental workflow. Alveolar macrophages (AM) and type II (AT2) cells were isolated from naïve (N) and influenza (F) treated young (3-4 months) (Y) and old (18 months) (O) mice by cell sorting. Proteins from the cell lysates were trypsin digested and subjected to 1D LC-MS/MS to identify peptides, which were subsequently mapped to their parent protein using MaxQuant and Andromeda software.

To characterize the identity of the detected proteins, we performed a preliminary categorization of all identified proteins for each cell type using the GO Slim categories of Biological Processes (**Table S1**) and Molecular Function (**Table S2**) available through PantherDB (56–58). This analysis revealed that our set of recovered proteins primarily populate GO Slim terms that include Cellular and Metabolic Processes with 34% and 28% coverage of terms respectively, followed by subcellular localization and biological regulation with a coverage of ∼14% of terms in these GO Slim categories (**Fig S2 & Table S1**). Additionally, the recovered proteins are primarily involved in cellular catalytic functions and protein binding with 32% and 30% of coverages of terms respectively. We did not observe any significant differences in the coverage of GO Slim terms between the 2 cell types, highlighting that the recovered proteins from AM and AT2 cells exhibit a similar distribution across biological processes (**Fig S2 & Table S1**) and molecular features (**Fig S3 & Table S2**) indicating that these cells are qualitatively similar despite their distinct biological roles and developmental specialization.

### Proteostasis network profiling of AM and AT2 cells

We were interested in the abundance of <u>p</u>roteostasis <u>n</u>etwork (PN) components in the proteomic data because their expression and function have a well-established role in protecting cells against environmental stress, including acute influenza infection (6,20,21,23,26,27,31). Additionally, the expression of PN components have been shown to be affected by age and protein misfolding stress. For our analysis, we used 2 different classifications of PN components, a full profile PN (14, 59), which includes 2708 proteins distributed across 16 biological categories (**Fig 2A & 2B**), and the more focused chaperome PN, which includes 318 proteins distributed across 9 different categories (**Fig 2C & 2D**) (31). We recovered 45% of full profile PN components (AM = 1210/2708; AT2 = 1239/2708) and 55% of chaperome PN components (AM = 174/318; AT2 = 176/318) in both cell types (**Fig 2**). The recovery of full profile PN components represents ∼30% of all identified proteins in the WCP analysis of AM and AT2 cells, highlighting the abundance of many of these components and emphasizing the central role of chaperones in biological pathways that manage the cellular response to aging and influenza challenge. A more in depth analysis revealed cell type-, age- and influenza-specific expression changes in these PN components as outlined below.

**Figure 2.**
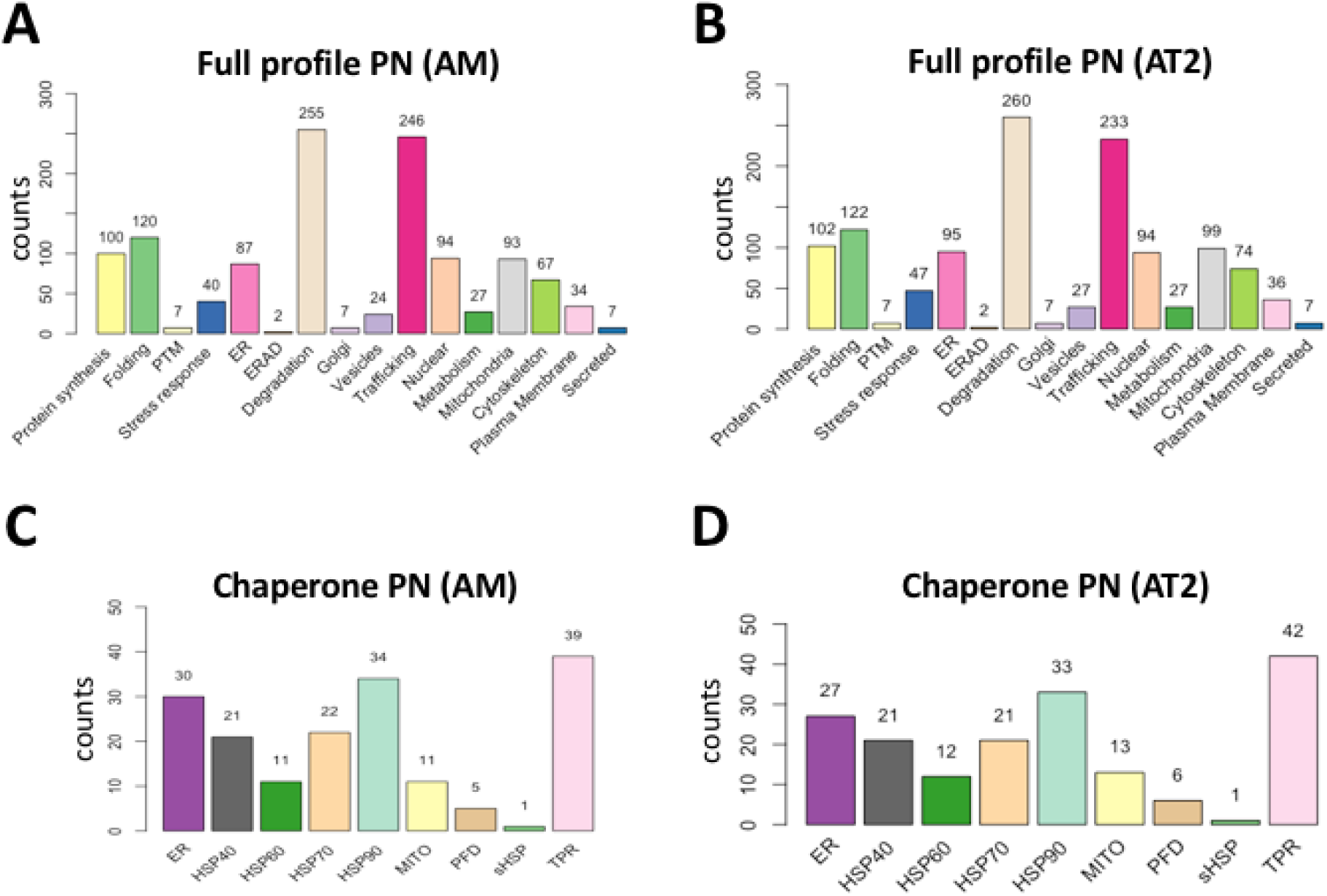
Alveolar cells exhibit similar proteostatic proteomic profiles. Bar graphs depicting the number of identified full profile proteostasis network (PN) proteins (14, 59) in AM (**A**) and AT2 (**B**) cells as well as chaperone PN proteins (31) in AM (**C**) and AT2 (**D**) cells. The number of identified proteins for each subcategory is indicated above each bar. The data represents proteins identified in at least 2 out of 3 replicates for at least one of the experimental conditions.

### Differentially abundant (DA) proteins in naïve and influenza challenged young and old cells

To understand the nature of the proteomic changes that account for the age-associated decline in cell health and tolerance to viral infection, we analyzed the WCP data sets from each treatment condition within the 2 cell types to highlight differentially abundant (DA) proteins across multiple transitions including young naïve (YN) versus old naïve (ON); ON versus old influenza (OF), YN versus young influenza (YF) and YF versus OF (**Fig 3A-D**). Here, we used the sum of the spectral intensities for all peptides mapping to each protein to determine their absolute abundance in a given sample. These protein intensity values were subsequently averaged across the triplicates for each condition and statistical significances (P < 0.05) DA were determined. Our analysis identified 1825 total AM-associated proteins and 1547 total AT2-associated proteins that exhibited a statistically significant DA in at least one of the transitions described above (**Fig 3A-D**). A specific analysis of the recovery of full profile PN components revealed that ∼27-30% of all DA proteins in all transitions described (**Fig 3A-D**) are PN components (**Fig 3E-H**), indicating that none of the transitions represent significant changes in the overall number of PN components that are exhibiting differential expression.

**Figure 3.**
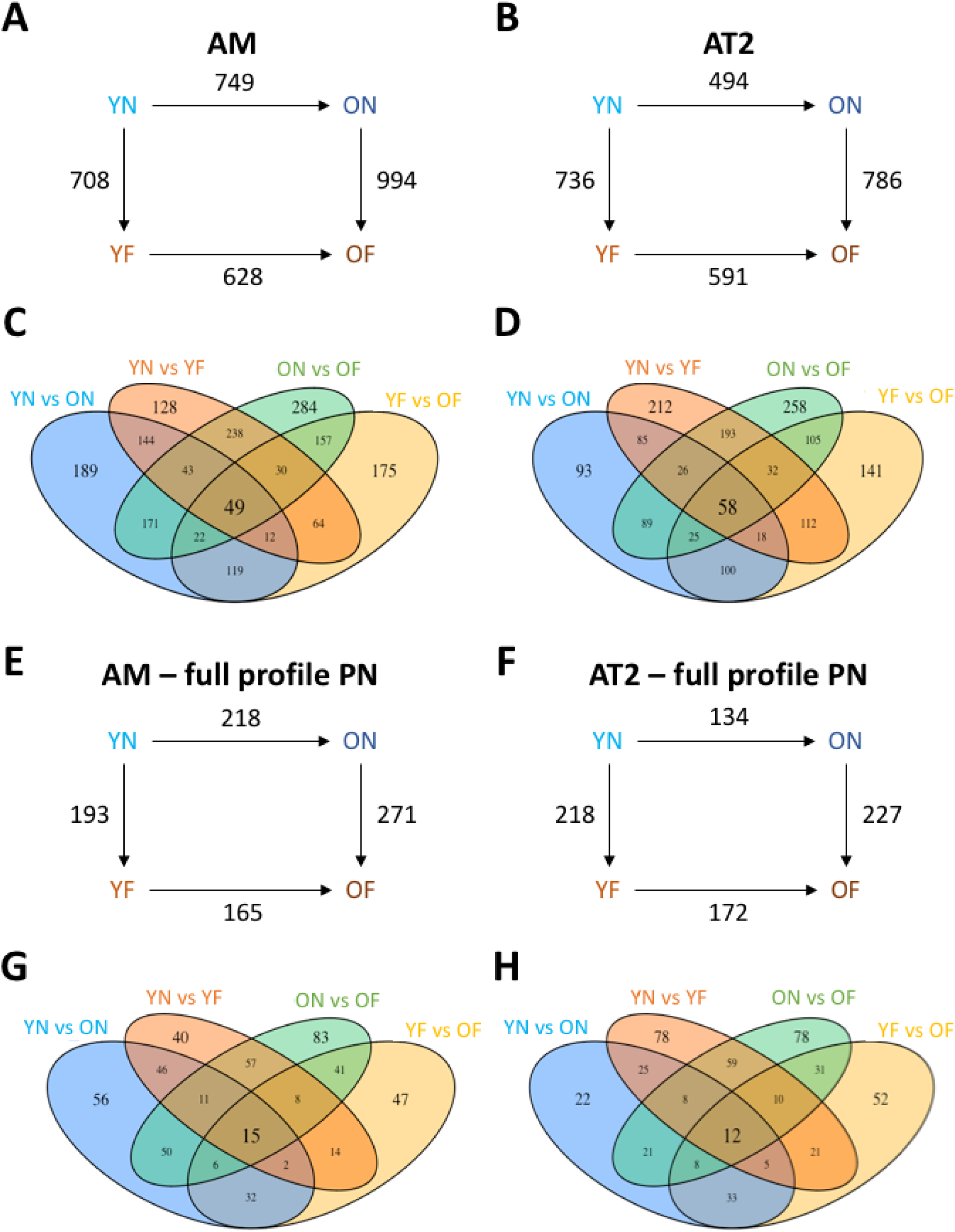
Alveolar cells exhibit unique patterns of differentially abundant proteins. Schematic diagram depicting the number of differentially abundant (DA) proteins (**A-D**) and proteostasis network (PN) components (**E-H**) between each of the depicted transitions, including young naïve (YN) to young influenza treated (YF); YN to old naïve (ON); ON to old influenza treated (OF) and ON to OF. Panels **A** and **B** depict the number of DA proteins between the indicated transitions for AM and AT2 cells respectively. Panels **C** and **D** depict the Venn diagram highlight the distribution of DA proteins between the various transitions in AM and AT2 cells respectively. Panels **E** and **F** depict the number of DA PN components between the indicated transitions for AM and AT2 cells respectively. Panels **G** and **H** depict the Venn diagram highlight the distribution of DA PN components between the various transitions in AM and AT2 cells respectively.

For AM-associated proteins we detected 994 DA proteins when comparing ON and OF, (**Fig 3A & 3C**), that is ∼30% higher than the comparison of YN and YF, where only 708 DA proteins were seen (**Fig 3A & 3C**). This difference is also reflected by the number of DA PN components, where 271 DA PN proteins are seen when comparing ON and OF compared to 193 DA PN seen when comparing YN and YF (**Fig 3E & 3G**). This reflects a differential response to influenza in old cells that either originates in significant differences in the response of the older cells to a viral challenge relative to younger cells, or a difference in the naïve proteomic profiles of young and old cells. Since the smallest number of DA proteins and PN components was seen in comparing YF and OF, where we observed 628 DA proteins (**Fig 3A & 3C**) and 165 DA PN components, respectively (**Fig 3E & 3G**), the differential response of older AM cells to influenza originates from large differences in the proteomic starting points of the O or Y naïve cells. Indeed, the second greatest number of DA proteins and PN components is seen when comparing YN to ON AM cells (**Fig 3A, 3C, 3E & 3G**). This trend is mirrored by the number of unique proteins and PN components for each transitions (**Fig 3C & 3G**), where we observe 284 unique DA proteins (**Fig 3C**) and 83 unique DA PN components (**Fig 3G**) in the pair-wise comparison of ON and OF, by far the largest category of unique statistically significant proteomic changes in our analysis of AM cells.

A different phenomenon is observed in AT2 cells. While we observe similar numbers of DA proteins and PN components when comparing the naïve to influenza infected cells between young and old cells, we observe the fewest number of DA proteins and PN components between the naïve states, where only 494 DA proteins (**Fig 3B**) and 134 DA PN components (**Fig 3F**) were detected. These results suggest that the naïve proteomes between young and old AT2 cells are more similar than for AM cells. In AM cells, we saw a decrease in the number of DA proteins after influenza challenge of YN and ON cells (**Fig 3A & 3C**) suggesting that despite their initial differences, their proteomes were getting closer to one another. In contrast, in AT2 cells, we see an increase in the number of DA proteins (**Fig 3B & 3D**) and PN components (**Fig 3E & 3H**) when comparing YF vs OF relative to that seen between YN and ON cells, suggesting that the YN and ON AT2 cells respond differently to this environmental stress.

An examination of whether DA PN components are upregulated or downregulated in naïve cells (**Fig 4, Fig S4-5**) revealed unique patterns of PN proteins that could contribute to the physiological effects of aging and response to influenza (**Fig 3**). For example, in AM cells (**Fig 4A, Table S1 & S2, Fig S4-5**), we observe a proteostatic collapse in aged cells, where a significant number of protein folding components are expressed at lower levels in older cells compared to younger cells, suggesting that aged AM cells have a reduced capacity to manage the influenza-mediated protein folding stress, which could require activation of stress response pathways. A similar profile is observed in AT2 cells suggesting that this decline reflects a systemic effect of aging (**Fig 4A & 4B - left, Table S1 & S2, Fig S4-5**). For example, we observe a disproportionate effect on all Hsp90-associated chaperome components in both AM and AT2 cells (**Fig 4A & 4B - right, Table S1 & S2, Fig S4-5**), as well as a significant decrease in Hsp70-associated chaperome components in AT2 cells (**Fig 4B – right, Table S1 & S2, Fig S4-5**) including tetratricoid<u>p</u>eptide repeat (TPR) containing chaperome components in AM cells (**Fig 4A – right, Table S1 & S2, Fig S4-5).**

**Figure 4.**
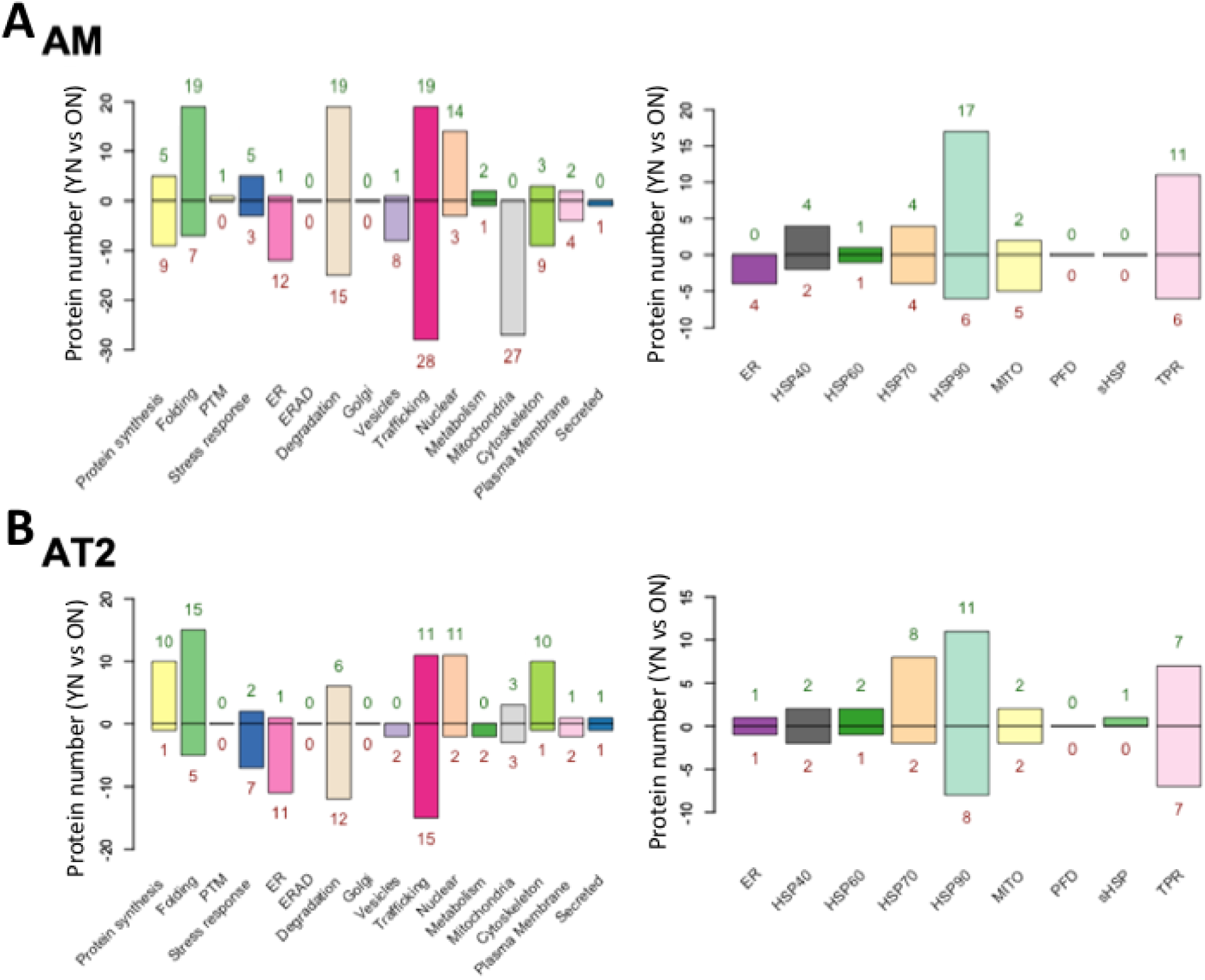
Alveolar cells exhibit unique expression patterns of proteostatic proteins. **A.** Bar graphs depicting the number of differentially abundant (DA) full profile (left) and chaperome (right) proteostasis network (PN) components between young naïve (YN) and old naïve (ON) AM cells. **B.** Bar graphs depicting the number of differentially abundant (DA) full profile (left) and chaperome (right) proteostasis network (PN) components between young naïve (YN) and old naïve (ON) AT2 cells. The number of DA proteins for each subcategory is indicated above each bar. The data represents a pairwise comparison of averaged protein intensity for each protein (n = 3) between YN and ON and those found to be statistically significantly different (p < 0.05) were indexed in this bar graph.

### Age-associated dysregulation of mitochondrial protein expression

Given that multiple pathways challenge chaperone management and hence the chronic aging process in AM and AT2 cells, and their response to acute influenza challenge, we performed a more extensive analysis of the biological processes that are the most affected by DA proteins. For this analysis, we first separated DA proteins from each cell type based on whether they were upregulated or downregulated in response to age and assessed which GO-associated pathways were the most affected by each of the protein sets (**Fig 5, FigS6-9**). For AM cells, we observed that proteins that are significantly upregulated with age are primarily associated with mitochondrial function and related proteostasis pathways (60–63), where 4 of the top 6 most impacted GO-associated pathways are linked to mitochondrial activity, including respiratory electron transport and the citric acid (TCA) cycle (**Fig 5A**). Conversely, for proteins in AM cells whose expression are significantly reduced with age, we observed that the most affected pathways include RNA localization and regulation, possibly related to maintenance of the transcriptome, as well as chaperome-mediated protein folding (**Fig 5B**). These data are consistent with our PN analysis of DA proteins where we observed increased mitochondrial protein expression and proteostatic ‘collapse’ with aging of AM cells (**Fig 4**). For AT2 cells, we observed that proteins upregulated with age primarily affect metabolic pathways, possibly related to mitochondrial function (32–37) (**Fig 5C**), whereas proteins that are downregulated with age primarily affect RNA stability and chaperone-mediated protein folding (**Fig 5D**), the latter being a similar observation to what was observed with downregulated proteins from AM cells (**Fig 5B**) and consistent with the observation of proteostatic collapse described in **Fig 4**.

**Figure 5.**
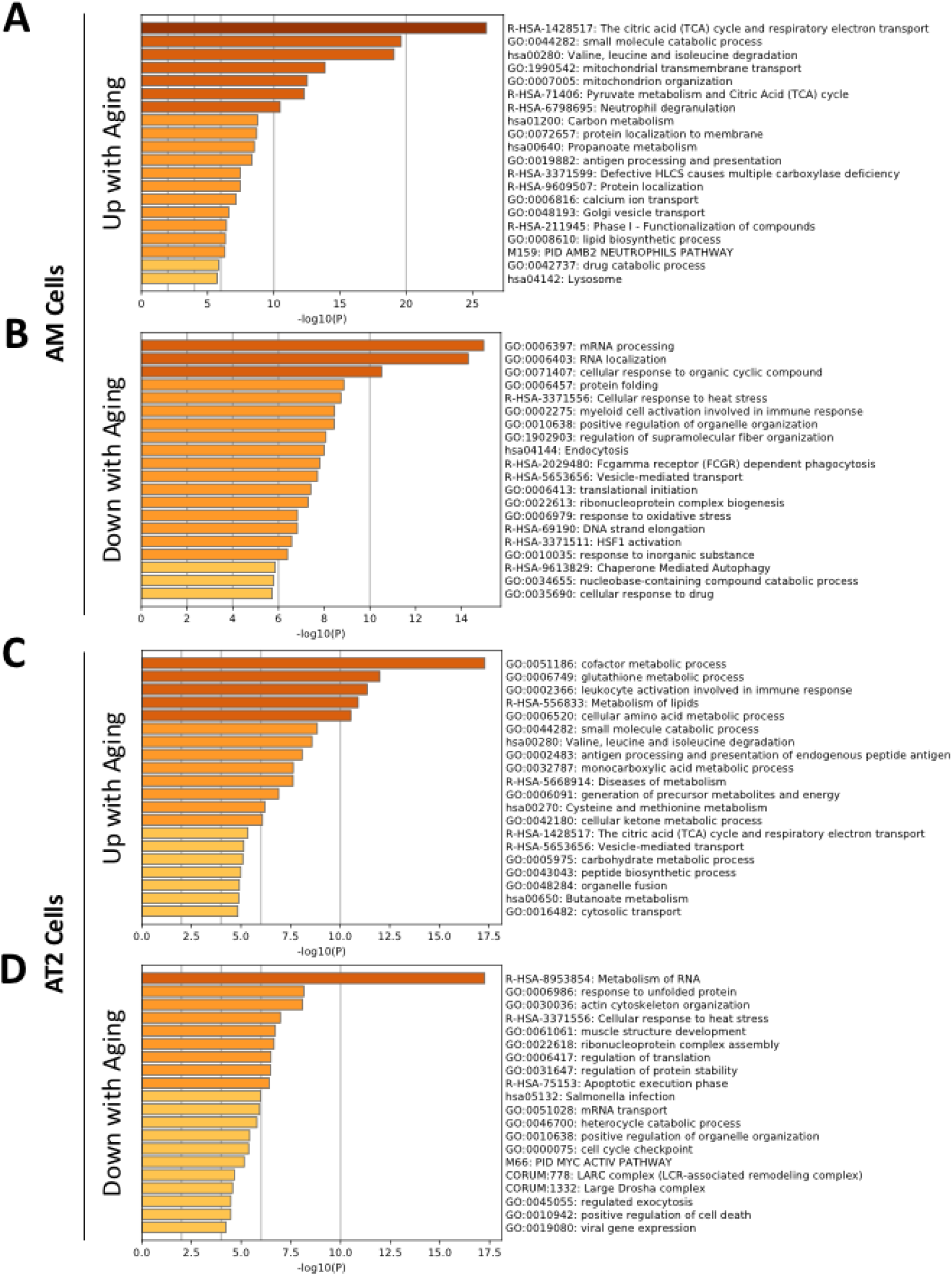
Differentially abundant proteins affect unique biological processes in different alveolar cells. **A.** Metascape analysis of biological processes impacted by the subset of proteins that are statistically significantly upregulated in alveolar macrophages (AM). **B.** Metascape analysis of biological processes impacted by the subset of proteins that are statistically significantly downregulated in alveolar macrophages (AM). **C.** Metascape analysis of biological processes impacted by the subset of proteins that are statistically significantly upregulated in alveolar type II cells (AT2). **D.** Metascape analysis of biological processes impacted by the subset of proteins that are statistically significantly downregulated in alveolar type II cells (AT2).

### Impact of mitochondria energetics on aging and influenza challenge

Given its central role in the management of energy reserves for ATP-intensive proteostasis function, we mapped the absolute expression of all mitochondrial proteins detected in our WCP analysis for all 4 conditions in both AM and AT2 cells (**Fig 6A**). An examination of these heat maps reveals a clear increase in mitochondrial proteins in aged AM cells relative to that seen in young AM cells (**Fig 6A – left**). We also observe differences in the expression of individual mitochondrial proteins in AT2 cells, and while the overall pattern of expression appears very similar we see a trend towards a decrease in individual mitochondrial protein expression with age as opposed to the increases seen in AM cells (**Fig 6A – right**). Moreover, the effect of influenza on AT2 cells resulted in an overall reduction in mitochondrial protein expression in both young and old cells (**fig 6A – right**). These results suggest that differences in the expression of mitochondrial proteins in AT2 cells do not account for the differential sensitivity of young and old cells to influenza. Conversely, while we see a trend towards a reduction in mitochondrial proteins in aged AM cells challenged with influenza (OF-AM), the overall expression pattern remains significantly higher than that seen in YF-AM cells (**Fig 6A – left**). These data suggest that the inability of old AM cells to reduce mitochondrial proteins sustains a higher demand for mitochondrial chaperones to manage the increased folding needs of this organelle. This stress like response could contribute to their hypersensitivity to influenza. Consistent with this interpretation, while in AM cells we see a striking increase in complex I components and complex III components (**Figure 6B, C - left**), AT2 cells show a more variable response with both increases and decreases reflecting the particular mitochondrial marker being assessed reflecting the ability of AT2 to manage the stress insult (**Fig 6B, C - right**). Thus, direct reads of proteome content suggest that individual cell types may differentially respond to mitochondria need and/or management in response to the challenges of chronic aging and acute stress insults. These results are consistent with the emerging importance of complex I and III on global cellular health during aging and its response through metabolic cascades including the TCA cycle (6,34–37,39,43,64,65).

**Figure 6.**
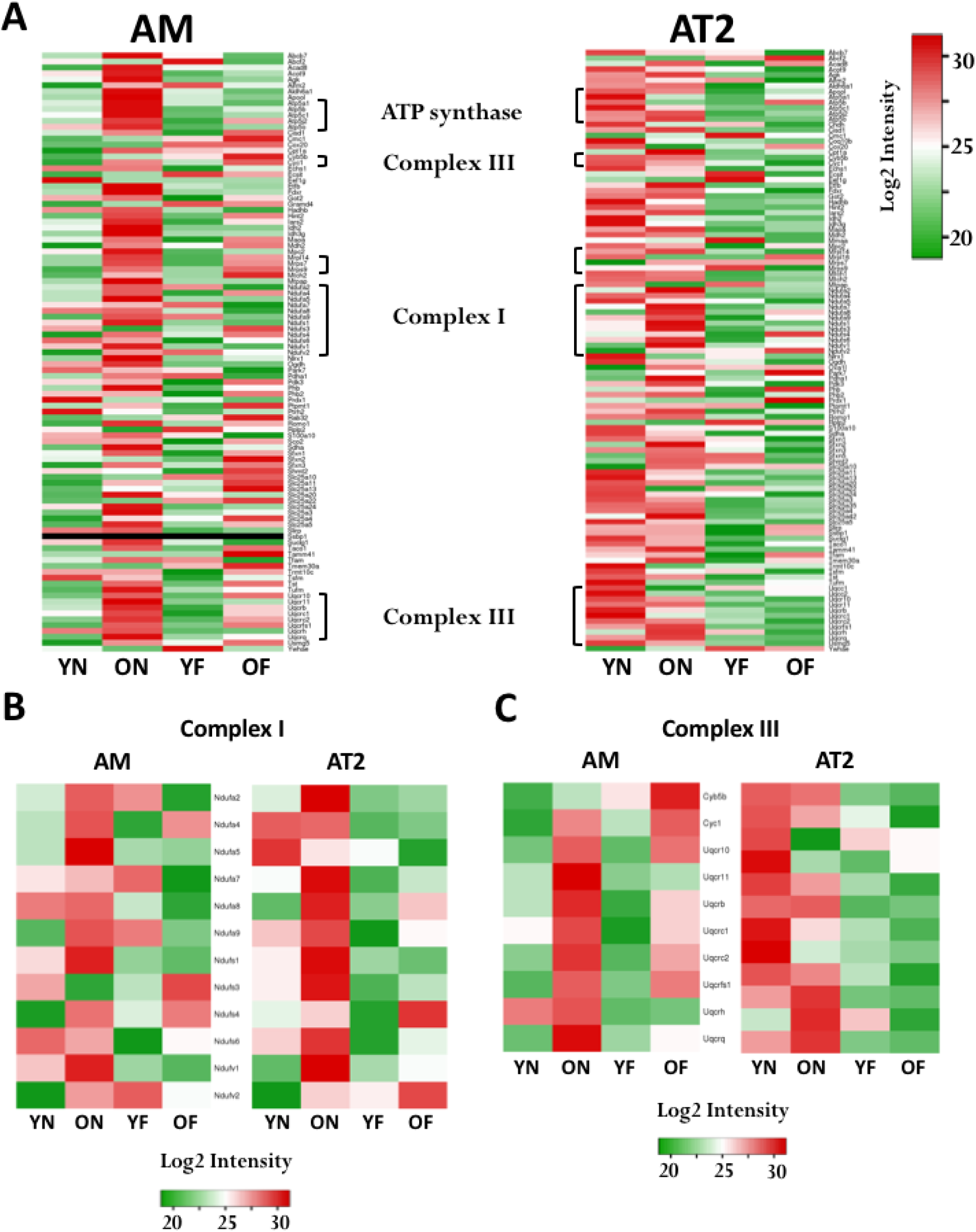
Aging causes a dysregulation of mitochondrial proteins in alveolar macrophages. **A.** Heat map depicting the log2 intensity of protein expression for all mitochondrial proteins identified in naïve (N) and influenza treated (F) young (Y) and old (O) alveolar macrophages (AM) (left) and alveolar type II (AT2) cells (right). **B.** Heat map depicting the log2 intensity of complex I mitochondrial proteins identified in naïve (N) and influenza treated (F) young (Y) and old (O) alveolar macrophages (AM) (left) and alveolar type II (AT2) cells (right). **C.** Heat map depicting the log2 intensity of complex III mitochondrial proteins identified in naïve (N) and influenza treated (F) young (Y) and old (O) alveolar macrophages (AM) (left) and alveolar type II (AT2) cells (right).

### Age-associated changes in proteostasis in influenza response

To further address the differential response of young and old mice to influenza challenge (**Fig 7A**), we divided our DA proteins into two groups. Group I represents DA proteins that are upregulated or unchanged in the YF vs YN transition but downregulated in the OF vs ON transition (**Fig 7B**). Group II represents DA proteins that are downregulated or unchanged in the YF vs YN transition and upregulated in the OF vs ON transition for each cell type (**Fig 7C**). This approach will highlight all DA proteins that exhibit differential response to influenza exposure (see Methods).

**Figure 7.**
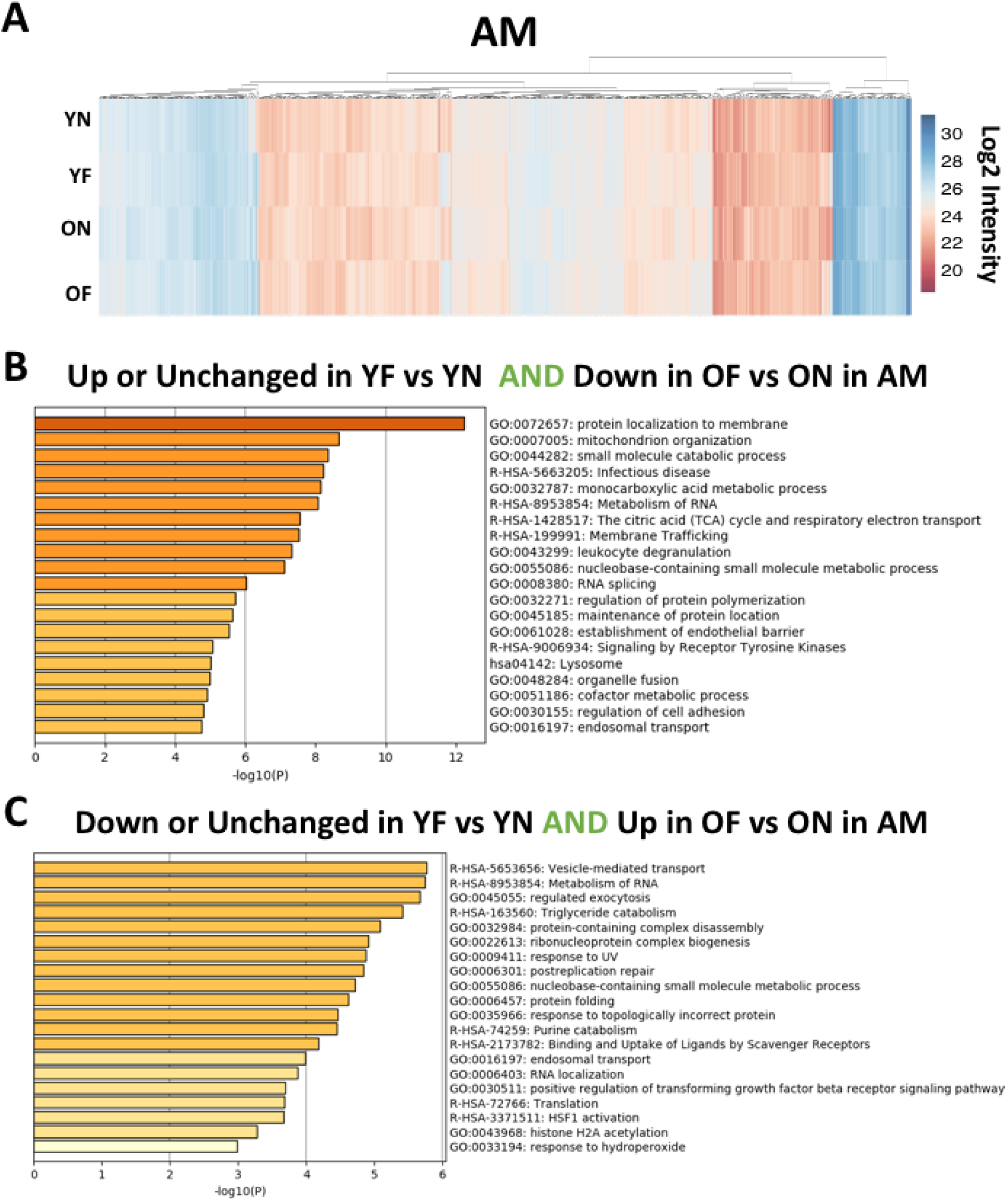
Viral infection causes differential proteomic patterns that map to distinct biological processes in alveolar macrophages. **A.** Heat map depicting the log2 intensity of all identified proteins in naïve (N) and influenza treated (F) young (Y) and old (O) alveolar macrophages (AM). **B.** Metascape analysis of biological processes impacted by proteins that are upregulated or unchanged in young influenza treated AM cells (YF) relative to their naïve state (YN) but downregulated in old influenza treated AM cells (OF) relative to their naïve state (ON). **C.** Metascape analysis of biological processes impacted by proteins that are downregulated or unchanged in young influenza treated AM cells (YF) relative to their naïve state (YN) but upregulated in old influenza treated AM cells (OF) relative to their naïve state (ON).

In AM cells, we identified 352 group I proteins and 260 group II proteins, highlighting that the response of ON cells to influenza challenge is strikingly different from YN cells (**Fig 7A & B**). Interestingly, a Metascape analysis of the group I and group II proteins sets revealed that they affect largely non-overlapping biological processes. For example, for group I proteins, we observe that young cells mount a response to infection, which is absent in aged AM cells (**Fig 7A & 7B**). Conversely, an analysis of group II proteins in AM cells reveals that aged cells have increased numbers of proteins involved in RNA metabolism, vesicle trafficking as well as protein translation and folding (**Fig 7A & 7C**), all processes that are typically abrogated in response to influenza stress, as seen in young AM cells. In AT2 cells we identified 287 group I proteins and 229 group II proteins. Again we observe that aged AT2 cells fail to mount a response to infection (**Fig 8A & 8B**), while continuing to sustain protein synthesis, folding and trafficking in the face of a mounting environmental stress (**Fig 8A & 8C**). These data suggest that aging lung cells fail to sense and/or respond to the viral stress and continue expressing biological systems that ultimately contributes to their demise.

**Figure 8.**
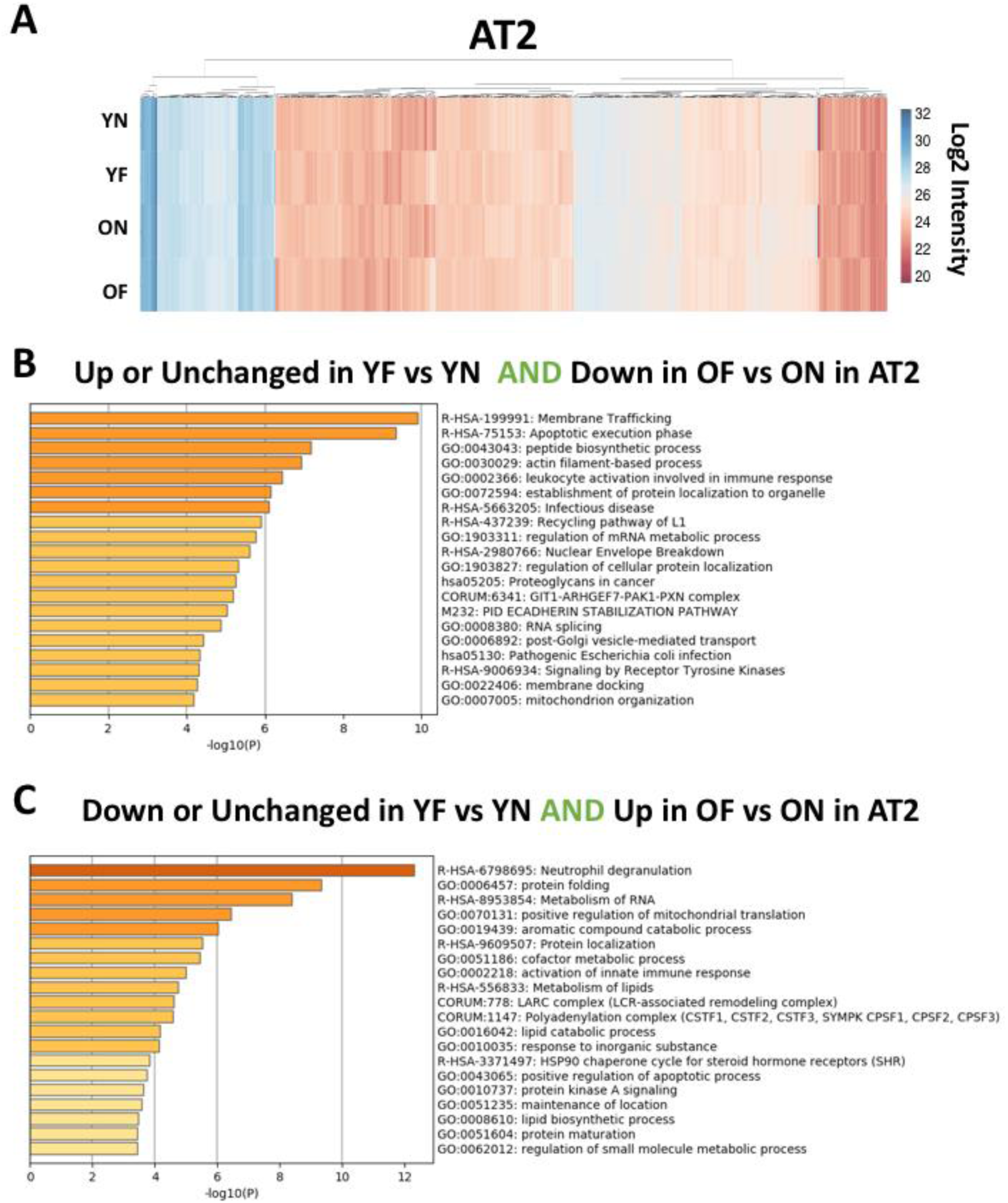
Viral infection causes differential proteomic patterns that map to distinct biological processes in alveolar type II cells. **A.** Heat map depicting the log2 intensity of all identified proteins in naïve (N) and influenza treated (F) young (Y) and old (O) alveolar type II cells (AT2). **B.** Metascape analysis of biological processes impacted by proteins that are upregulated or unchanged in young influenza treated AT2 cells (YF) relative to their naïve state (YN) but downregulated in old influenza treated AT2 cells (OF) relative to their naïve state (ON). **C.** Metascape analysis of biological processes impacted by proteins that are downregulated or unchanged in young influenza treated AT2 cells (YF) relative to their naïve state (YN) but upregulated in old influenza treated AT2 cells (OF) relative to their naïve state (ON).

To understand how the cellular PN components (**Fig 2-4**) are responding aging and viral stress, we turned our focus to characterize the expression of full profile PN proteins in response to the influenza challenge of YN and ON cells. A comparison of the DA proteins between the YN versus ON and YF versus OF transitions reveal that while there are some attempts by the old cells to establish a more youthful response to this environmental stress, the proteomes remain significantly different in both AM (**Fig 9A**) and AT2 (**Fig 9B**) cells. Specifically, we observe that the PN profile of folding components between Y vs O cells after influenza challenge exhibit a more similar profile than observed in the naïve state (**Fig 9A**). This effect is due to both a reduction in folding proteins in YF AM cells as well as an increase in folding proteins in OF AM cells. We also observed a more similar profile for mitochondrial PN proteins between Y and O AM cells relative to that seen in their naïve states (**Fig 9A**), an effect that is due to a reduction in mitochondrial proteins (**Fig 6A**) after viral challenge of aged AM cells, including a reduction of complex I (**Fig 6B**) and complex III components (**Fig 6C**). In the case of AT2 cells, we observed a more similar PN profile for protein translation and folding components after viral challenge of both Y and O AT2 cells relative to that seen in their naïve states (**Fig 9B**). These effects are due to increased expression of protein translation and folding components in OF AT2 cells as well as reduction of protein folding proteins in YF AT2 cells. The need for OF AM and AT2 cells to activate stress response pathways to accommodate their increased need for chaperome proteins could represent a significant obstacle to their ability to survive this viral infection. The activation of stress response pathways is often accompanied by abrogation of global protein synthesis, which could curb the expression of other key cellular proteins needed for immediate and long term management of cellular processes. We also observed that the PN profile for folding, degradation, trafficking and cytoskeletal PN components became more divergent than they initially were in their naïve states (**Fig 9B**), a phenomenon not observed for any PN categories in AM cells. These results suggest a divergence in the management of protein folding stress in response to both aging and influenza challenge in AM and AT2 cells.

**Figure 9.**
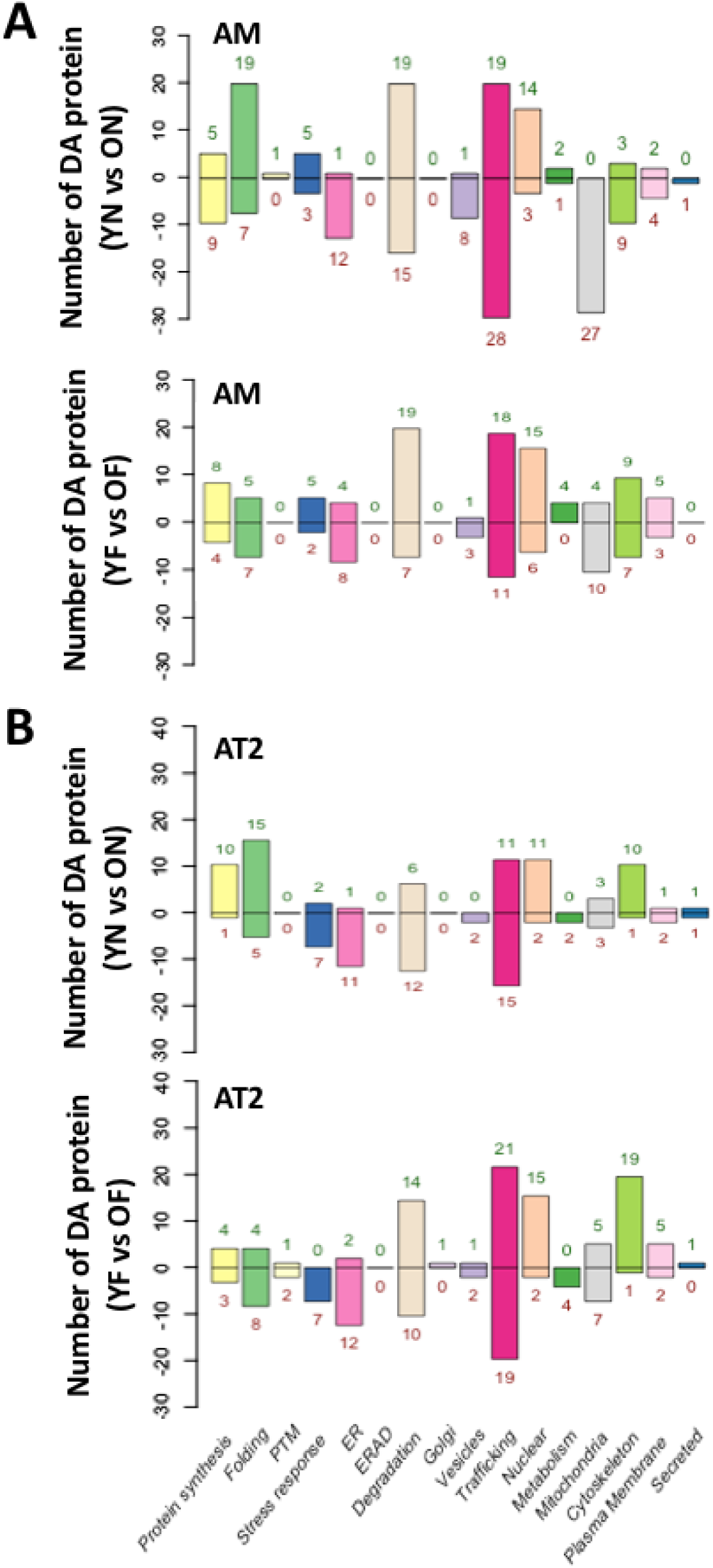
Influenza A infection induces unique proteostatic patterns of expression in alveolar cells. **A.** Bar graphs depicting the number of differentially abundant (DA) full profile proteostasis network (PN) components between young naïve (YN) and old naïve (ON) transition (top) and young influenza treated (YF) and old influenza treated (OF) (bottom) AM cells. **B.** Bar graphs depicting the number of differentially abundant (DA) full profile proteostasis network (PN) components between young naïve (YN) and old naïve (ON) transition (top) and young influenza treated (YF) and old influenza treated (OF) (bottom) AT2 cells. The number of DA proteins for each subcategory is indicated above each bar. The data represents a pairwise comparison of averaged protein intensity for each protein (n = 3) between the indicated transitions and those found to be statistically significantly different (p < 0.05) were indexed in this bar graph.

### Folding components differentially regulated by aging and influenza challenge

To begin to understand which folding chaperones specifically contribute to the differential response of young and aged lung cells to pathogenic challenge by influenza, we generated chaperone network maps to highlight the differential changes in young (YN) and old (ON) cells and in response to influenza (YF & OF) to delineate changes that would influence client protein synthesis, folding and stability unique to each cell type and aged state (**Fig 10A, upper panel, (YN-ON, YN-YF, ON-OF, and YF-OF))**. The differential chaperone network for each transition illustrates the percent increase (closed circles) and decrease (open circles) in the expression of each folding chaperone component and exemplifies how the client folding network in AM cells (**Fig. 10A**) and AT2 cells (see below, **Fig. 10B**) is being reprogramed with age and by influenza challenge.

**Figure 10.**
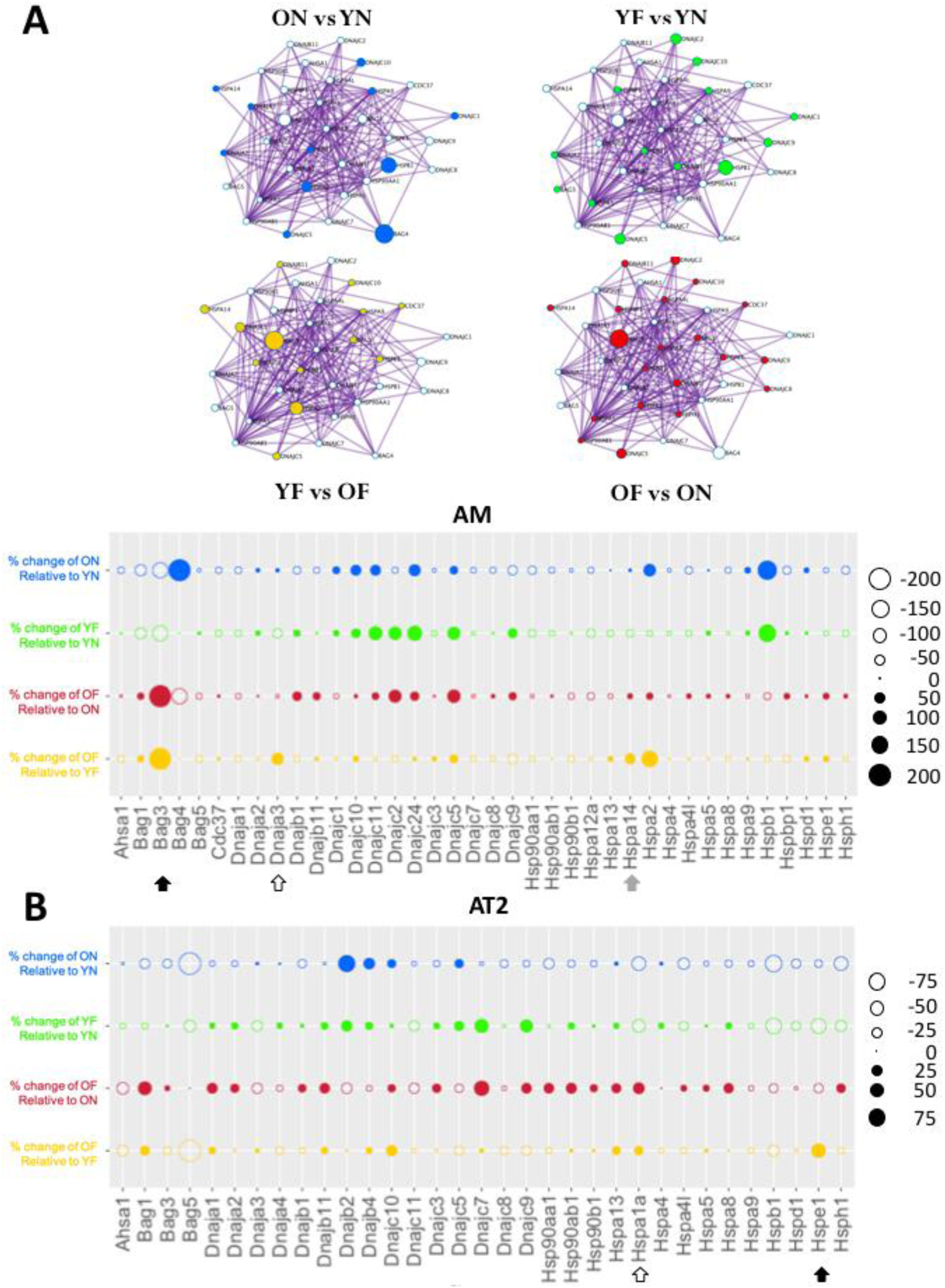
Influenza infection induces unique patterns of folding chaperone expression in alveolar macrophages. **A. (upper panel)** Network analysis of chaperone-PN folding components for ON vs YN (blue), YF vs YN (green), OF vs ON (red) and YN vs ON (yellow). **(lower panel).** Dot plot representing the percent change in the expression of the indicated folding chaperone PN components for the indicated transitions in alveolar macrophages (AM), which include young naïve (YN) to old (naïve) (blue); YN to young influenza treated (YF) (green); old naïve (ON) to old influenza treated (OF) (red) and YF to OF (yellow). **B.** Dot plot representing the percent change in the expression of the indicated folding chaperone PN components for the indicated transitions in alveolar type II cells (AT2), which include young naïve (YN) to old (naïve) (blue); YN to young influenza treated (YF) (green); old naïve (ON) to old influenza treated (OF) (red) and YF to OF (yellow). The circle size represents the magnitude of the percent change in the indicated transition. The closed circles represent positive percent changes, whereas open circles represent negative percent changes.

To more easily depict the changes that occur in each of these transitions in AM cells, the data were replotted in a network independent manner (**Fig. 10A, lower panel**). While modest changes were observed in core Hsp70 and Hsp90 chaperones, the data highlight that the major changes affecting aging and the age-dependent response to influenza are found in the co-chaperones impacting chaperone function, referred to as the CARE network. We define the CARE network as a subset of the components contributing to the UPR, the ISR-UPR_mito_ pathway, and the HSR proteostasis signaling pathways, the latter charged with managing client delivery to the cytosolic core Hsp70 and Hsp90 chaperones that control protein folding and stress response to misfolding. Interestingly, changes in CARE network components contribute to the management of Hsp70 and Hsp90 orthologs in the multiple compartments including the cytosol, the ER and mitochondria (**Fig. 10A, lower panel**).

For example, we observed significant changes in the expression of Bcl-2 associated athanogene (Bag) proteins, which increase client release from Hsp70 by accelerating nucleotide exchange at the ATP site of this folding chaperone, in both naïve (ON) and influenza challenged (OF) aged cells. Here, we observed a striking difference in the expression pattern of Bag3 (**Fig 10A, lower panel, black arrow**), which is involved in regulating autophagy pathways (14,66–77), in response to influenza. The expression of Bag3 was higher in YN-AM cells relative to that seen in ON-AM cells (**Fig 10A, lower panel, black arrow – blue circle**), however, upon influenza challenge, we observed that young AM cells decreased Bag3 expression (**Fig 10A, lower panel, black arrow – green circle**), whereas old AM cells increased Bag3 expression (**Fig 10A, lower panel, black arrow – red circle**) – resulting in a dramatically different chaperome response between YF and OF (**Fig 10A, lower panel, black arrow – yellow circle**).

We also noted a significant differences with the mitochondrial chaperone protein, Hspa14 (Hsp60) (78, 79) (**Fig 10A, lower panel, grey arrow**) and its mitochondrial associated co-chaperone DnaJa3 (80) (**Fig 10A, lower panel, white arrow**) to influenza exposure, both of which exhibit a marginally higher basal expression in old naïve cells (**Fig 10A, lower panel, grey and white arrows - blue**). These results are consistent with the increased expression of mitochondrial proteins in ON-AM cells (**Fig 6A**). Whereas Hsp60 exhibits a modest increased expression in older ON-AM relative to YN-AM cells (**Fig 10A, lower panel, grey arrow – blue circle**), we observe that exposure of young AM cells to influenza decreases the expression of this chaperone (**Fig 10A, lower panel, grey arrow – green circle**), whereas the same dose of influenza causes older AM cells to increase expression (**Fig 10A, lower panel, grey arrow – red circle**), resulting in a dramatically different response between young and old AM cells and corresponding co-chaperone managers (**Fig 10A, lower panel, grey arrow – yellow circle**). Interestingly, the response of both young and old mice to influenza is to reduce the expression of DnaJA3 (**Fig 10A, lower panel, white arrow – green & red circles**), a task for which young AM cells are more effective than older AM cells (**Fig 10A, lower panel, white arrow – green circle**), resulting in a significantly different response to influenza exposure (**Fig 10A, lower panel, white arrow – yellow circle**).

In the case of the AT2 cells, we observe 2 primary differences in the responses of young and old cells to influenza as shown in a network independent manner (**Fig 10B**). The first we observe is in the response of Hspe1 (Hsp10), a mitochondrial small heat shock protein (sHsp) co-chaperone involved in Hsp70-dependent mitochondrial protein import, which is more highly expressed in YN relative to ON cells (**Fig 10B, black arrow, – blue circles**). The influenza challenge causes Hspe1 to be downregulated in both young and old AT2 cells (**Fig 10B, black arrow – green & red circles**), but the old cells appear less capable of mounting the same response to that seen in the young AT2 cells resulting in a significantly elevated expression in OF-AT2 cells relative to that seen in YF-AT2 cells (**Fig 10B, black arrow – yellow circles**), leading to proteostasis collapse. The second folding chaperone for which we see a differential response to influenza between young and old AT2 cells is with the expression of HspA1A (Hsp70), whose activity is affected by the CARE Bag and DnaJ family co-chaperones. A comparison of YN- vs ON-AT2 cells reveals that HspA1A is more highly expressed in YN cells (**Fig 10B, white arrow – blue circle**), consistent with our observation above that old AT2 cells exhibit a proteostatic collapse (**Fig 4B**), a phenomenon also highlighted by the general upregulation (**Fig 10B- open circles**) of folding chaperones in YN-AT2 cells relative to that seen in ON-AT2 cells (**Fig 10B – blue**). The exposure to influenza results in a downregulation of HspA1A in young AT2 cells but upregulation in old AT2 cells (**Fig 10B, white arrow – green & red circles respectively**). The response of older cells therefore appears to be a dysregulated response to virus infection since OF-AT2 cells express more HspA1A than that seen in YF-AT2 cells (**Fig 10B, white arrow – yellow circles**). This HSR-like activation would typically be accompanied by a cell-wide reductions in protein expression, given the known maladaptive response of the HSR to chronic misfolding in rare disease (16), which presents a significant barrier to the recovery of these older cells to viral infection.

Although we observe changes in the expression of core Hsp70 and Hsp90 components, the majority of the changes appear to stem from changes in the regulatory co-chaperones. These data highlight that a potentially important driver of both aging and influenza-related pathophysiology are empowered by the poorly understood CARE contingent of the UPR, ISR-UPR_mito_, and HSR PN signaling pathways, which fails in aging, likely reflecting a chronic, possibly maladaptive, challenge to fold maintenance(16). This proteostatic challenge is further exacerbated by acute environmental challenges such as influenza leading to activation of stress response pathways in proteostatically collapsed aged cells contributing to cell death, a conclusion consistent with the projected global role of proteostasis response pathways managing human health and disease throughout a lifespan (6,17,81).

## Discussion

We have presented the results of a comprehensive proteomic analysis of sorted AM and AT2 cells from the lungs of naïve and influenza compromised young and old mice. These are key cell populations contributing to the overall function and protection of the lung over a lifespan. In general, we observe changes that reflect a challenge to the function of the prevailing PN in ER, cytosolic and mitochondrial pathways managing cellular energetics in response to protein misfolding burden in aged AM and AT2 proteomes. These changes could provide insight into why older mice are acutely sensitive to influenza. The observed attempt of aged cells to mount a chaperone-mediated defense to manage the viral challenge, places these cells in a state of maladaptive stress (16) which would be accompanied by significant global changes in the proteome stability, compromising their ability to manage the progression of and/or recovery from influenza infection and subsequent pneumonia-related stress challenges (9).

Although GO annotation reveals multiple adjustments to the proteome over the lifespan, two themes of contemporary relevance focused our attention on a subset of changes, namely mitochondrial related pathways (6,34–37,39,43,64,65) and proteostasis pathways (18–28). Specifically, in mitochondrial pathways and related mitochondrial stress responsive pathways (60–63), changes that are now implicated in the aging process from multiple perspectives. In particular, both complex I and complex III and their links to key metabolic pathways are asserted to be drivers of the aging process (6,34–37,39,43,64,65). Herein, we noted substantial changes in the expression of mitochondrial proteins involved in both complexes, with a marked increase in abundance of mitochondrial components in AM cells compared to AT2 cells, suggestive of cell specific differential responses reflecting their unique roles in the pathobiology of the aging process and differential response to chronic particulate and pathogen environmental stresses such as viral/bacterial challenge leading to pneumonia in the elderly and long-term impact on recovery (9). The latter results raise the possibility that it is not mitochondrial loss per se in some cell types (82), but rather a mitochondrial maladaptive proteomic response reflecting the need to sustain energy production that places an excessive burden on the mitochondrial proteostasis, an effect that is further exacerbated when these aged cells are exposed to an external stress, such as acute influenza infection.

Additionally, we observed changes in the expression of multiple proteostasis components involved in the functionality of the Hsp70 and Hsp90 core chaperome complexes, which have previously been linked to the management of neurodegenerative and other protein misfolding challenges during aging (18–28). The specific changes we observed herein focused our attention on the co-chaperone components (CARE) managing Hsp70 control of client folding as a central player in aging. In particular, the Bag3 response, being one of the largest differential responses detected, is involved in autophagic pathways, suggests that core chaperone associated degradation pathways associated with autophagy may be central to the management of AM and AT2 cells during aging, as suggested previously for other aging-related aggregation prone states (18–28). These observations raise the possibility that aging of the lung, a major contributor to the decline of lung function in response to environmental stress over a lifetime (9), is, at least in part, an uncharacterized aggregation prone disease, perhaps in response to supersaturation (83, 84) that must be precisely managed by the extensive Bag and DnaJ families through CARE over a lifespan. We also noted that while the Hsp90 core chaperone only exhibited modest changes in abundance (**Fig 10**), the Hsp90 chaperone/co-chaperone system as a whole underwent large differential changes in both AM and AT2 cells (**Fig 4**), suggesting that, like observed for Hsp70, it is the CARE network components that dictate the success of the chaperone network to manage the misfolding challenges that arise with aging. Considerable experimental efforts will need to be devoted to understanding the specificity of the collective role of CAREs in the maintenance and repair pathways contributing to human healthspan.

Not unlike changes observed in naïve young vs old AM and AT2 cells, influenza challenge yielded differential effects on both young and old cell types. Again, GO annotation reveals changes in multiple categories, but the cumulative evidence suggests that the PN, particularly the co-chaperone subnetworks controlling the chaperoning ability of Hsp70 orthologs, play critical roles in the response to and management after viral exposure (85, 86). Given that the proteomic analysis of influenza response revealed the potential importance of select Bag family proteins, DnaJ/Hsp40 family members and sHsp proteins, we suggest that the kinetics of client binding and release from the Hsp70 system could be a central regulator of influenza pathophysiology of the lung contributing to the age-related proteostasis collapse and maladaptive stress response mediated repair in the elderly that could be adjusted therapeutically.

A major challenge for understanding the utility of high value, comprehensive proteomic datasets is to understand the basis for the overall proteomic response in health and disease, and how these responses are integrated in different ways in different individuals to manage lifespan in a fashion that makes each one of us unique (47,49,87,88). We have proposed the principle of spatial covariance (SCV) (19) whereby spatio-temporal adjustments to the protein fold and/or composition of the folding PN can be captured by applying <u>G</u>aussian <u>p</u>rocess regression (GPR) based <u>m</u>achine learning (GPR-ML) (19). We have demonstrated that GPR-ML provides a computational platform to universally address the misfolding challenges facing the population in the context of individuals harboring disease-causing mutations during their lifespan. It is now apparent that the principle of SCV, and related machine learning approaches (50), will be essential to understand the role of the client and their associated PN network in managing protein folding in response to aging and to environmental stress related challenges (9). SCV principles will also be critical to understand the functionality of the proteome in the context of genome design and transcriptome responses that generate the protective barriers (3, 35) and CARE, as shown herein, that limit or attempt to limit the impact of aging- and pathology-associated stress to achieve a more sustainable level of lung function over a lifespan (13,89–92).

## METHODS

### Animals

All mouse procedures were approved by the Institutional Animal Care and Use Committee at Northwestern University (Chicago, IL, USA). All strains including wild-type mice are bred and housed at a barrier and specific pathogen–free facility at the Center for Comparative Medicine at Northwestern University. Male C57BL/6 mice were provided by NIA NIH and were housed at Northwestern University Feinberg School of Medicine Center for Comparative Medicine for 4 weeks prior to sacrifice. Mice were euthanized by pentobarbital sodium overdose. Immediately the chest cavity was opened and animals were perfused via the left ventricle with 10 mL of HBSS (Ca/Mn+). The lungs were harvested and subjected to enzymatic digestion to release stromal and immune cells and sorted by fluorescence-activated cell sorting. Cells were washed 3 times with ice cold PBS to remove all traces of blood proteins associated with the cells and FBS found in the collection media. Cells were subsequently frozen on dry ice and stored at −80°C until processing.

### Influenza A virus infection

Influenza virus strain A/WSN/1933 (WSN) was grown for 48 h at 37.5°C and 50% humidity in the allantoic cavities of 10–11-d-old fertile chicken eggs (Sunnyside Hatchery, WI). Viral titers were measured by plaque assay in Madin-Darby canine kidney epithelial cells (American Type Culture Collection). Virus aliquots were stored in liquid nitrogen and freeze/thaw cycles were avoided. For infection, mice were anesthetized with isoflurane and infected intratracheally with 10 plaque forming units (pfu) in 50 μL of PBS. Mice were sacrificed after 4 days.

### Tissue preparation and flow cytometry

Tissue preparation for cell sorting was performed as previously described (55, 65). Sample processing, washing and sorting was performed in protein-free buffers. For single-cell suspension obtained from tissues, erythrocytes were lysed using BD Pharm Lyse, and cells were counted using Nexcelom K2 Cellometer C automated cell counter (Nexcelom) with AO/PI reagent. Cells were stained with eFluor 506 (eBioscience) viability dyes, incubated with FcBlock (BD), and stained with fluorochrome-conjugated antibodies (see STAR Methods). Data and cell sorting were performed at Northwestern University RLHCCC Flow Cytometry core facility on SORP FACSAria III, BD LSR II, BD Fortessa and BD Symphony instruments (Becton Dickinson). Sorting was performed using a 100-µm nozzle and 40 psi pressure. Compensation, analysis and visualization of the flow cytometry data were performed using FlowJo software (Tree Star). “Fluorescence minus one” controls were used when necessary to set up gates.

### Sample preparation for proteomic analysis

Cells lysates were prepared by adding lysis buffer (8 M urea, 50 mM ammonium bicarbonate buffer) to thawed cells and subsequently digesting with trypsin. Briefly, cysteine disulfide bonds were reduced with 5 mM Tris(2-carboxyethyl) phosphine (TCEP) at 30 °C for 60 min, followed by cysteine alkylation with 15 mM iodoacetamide (IAA) in the dark, at room temperature for 30 min. Following alkylation, urea was diluted to 1 M urea using 50 mM ammonium bicarbonate, and proteins were finally subjected to overnight digestion with mass spec grade Trypsin/Lys-C mix (Promega, Madison, WI). The digested proteins were desalted using AssayMap C18 cartridges mounted on a BRAVO liquid handling system (Agilent, Columbia, MD), and the organic solvent was removed in a SpeedVac concentrator prior to LC-MS/MS analysis.

### LC-MS/MS procedures

Dried samples were reconstituted with 2% acetonitrile, 0.1% formic acid and analyzed by 1D-LC-MS/MS using a Proxeon EASY nanoLC system (Thermo Fisher Scientific) coupled to an Orbitrap Fusion Lumos mass spectrometer (Thermo Fisher Scientific). Peptides were separated using an analytical C18 Acclaim PepMap column 0.075 × 500 mm, 2 μm particles (Thermo Scientific) in a 90-min linear gradient of 2–28% solvent B at a flow rate of 300 nL/min. The mass spectrometer was operated in positive data-dependent acquisition mode. MS1 spectra were measured with a resolution of 120,000, an AGC target of 1e6, a maximum injection time of 100 ms and a mass range from 350 to 1400 m/z. The instrument was set to run at top speed mode with 3 s cycles for the survey and the MS/MS scans. After a survey scan, tandem MS was performed on the most abundant precursors exhibiting a charge state from 2 to 8 of greater than 5e3 intensity by isolating them in the quadrupole at 0.8 Th. HCD fragmentation was applied with 30% collision energy and the resulting fragments were detected using the turbo scan rate of the ion trap. The AGC target for MS/MS was set to 1e4 and the maximum injection time limited to 15 ms. The dynamic exclusion was set to 15 s with a 10 ppm mass tolerance around the precursor and its isotopes.

### LC-MS/MS data analysis

All mass spectra were analyzed with MaxQuant software version 1.5.5.1 (93). MS/MS spectra were searched against the Homo sapiens Uniprot protein sequence database (version July 2017) and GPM cRAP sequences (commonly known protein contaminants). Precursor mass tolerance was set to 20 ppm and 4.5 ppm for the first search where initial mass recalibration was completed and for the main search, respectively. Product ions were searched with a mass tolerance 0.5 Da. The maximum precursor ion charge state used for searching was 7. Carbamidomethylation of cysteines was searched as a fixed modification, while oxidation of methionines and acetylation of protein N-terminal were searched as variable modifications. Enzyme was set to trypsin in a specific mode and a maximum of two missed cleavages was allowed for searching. The target-decoy-based false discovery rate (FDR) filter for spectrum and protein identification was set to 1%.

### Statistical analysis

The evidence table output from MaxQuant was used for label-free protein quantitative analysis. First, calculated peptide intensities were log2-transformed and normalized across samples to account for systematic errors. A total of 8 normalization approaches were deployed (Loess, Robust Linear Regression, Variance Stabilization and Normalization, Total Intensity, Median Intensity, Average Intensity, NormFinder and Quantile), and their performance assessed (94) in order to determine the optimal normalization method (herein Loess normalization). Following normalization, all non-razor peptide sequences were removed from the list. Protein-level quantification and testing for differential abundance were performed using MSstats bioconductor package (95, 96) based on a linear mixed-effects model. The model decomposes log-intensities into the effects of technical and biological replicates, peptides and statistical interactions.

### Bioinformatic analysis

The initial list was filtered to remove protein contaminants and proteins not detected in at least two of three replicates from each experimental group. For each protein in each of the groups, the log2 median of the replicated intensities was considered. Initial GO Slim categorization was performed using the GO-Slim tool available through the Panther database (56–58), using GO Slim categories from Biological Process and Molecular function. Proteostasis annotation was obtained from two sources: 1) Brehme et al. (31) for chaperome annotation, and 2) a manually curated collection of over 2500 proteostasis components (14), where previous partial lists were merged using the Protein Atlas (97) as primary source for subcellular localization, protein function and proteostasis classification. Such annotation is available at https://github.com/balchlab/PPT_annotation.Differentially abundant proteins between experimental groups were selected according to a p-value of ≤0.05 as inclusion threshold.

To address differences in the how young and old mice respond to Flu, we first computed the delta between Naïve and Flu Intensities for both young and old (ΔY and ΔO). We obtain:

> ΔY ≥ 0 : proteins in Young that are up regulated or unaffected by Flu;
>
> ΔY < 0 : proteins in Young that are down regulated by Flu;
>
> ΔO ≥ 0 : proteins in Old that are up regulated or unaffected by Flu;
>
> ΔO < 0 : proteins in Old that are down regulated by Flu;

Hence we focus on the following subsets:

> Up/Unchanged in Young+F (ΔY ≥ 0) AND Down in Old+F (ΔO < 0).
>
> Down in Young+F (ΔY < 0) AND Up/Unchanged in Old+F (ΔO ≥ 0).

Gene Ontology (GO) and pathway membership analysis from the lists of differentially regulated proteins were obtained using Metascape (98). KEGG, Reactome and GO database results are presented as enriched GO term bar graphs, color-coded from grey to brown according to log10(p) of the standard cumulative hypergeometric statistical test. R/Bioconductor was used for data manipulation, analysis and visualization. Annotations and mappings were obtained from Uniprot (3/2019) and org.Mm.eg.db v.3.8.2 (Bioconductor release 3.9). Cytoscape (99) was used for interactive network visualization and presentation.

## Acknowledgements

WEB supported by NIH AG049665, HL095524, DK051870, HL141810; GRSB supported by GRSB: NIH AG049665. K.R.: NIH HL079190 and HL124664. RIM: NIH AG049665. Northwestern University Flow Cytometry Facility is supported by NCI Cancer Center Support Grant P30 CA060553 awarded to the Robert H. Lurie Comprehensive Cancer Center.

---

**Table 1.**
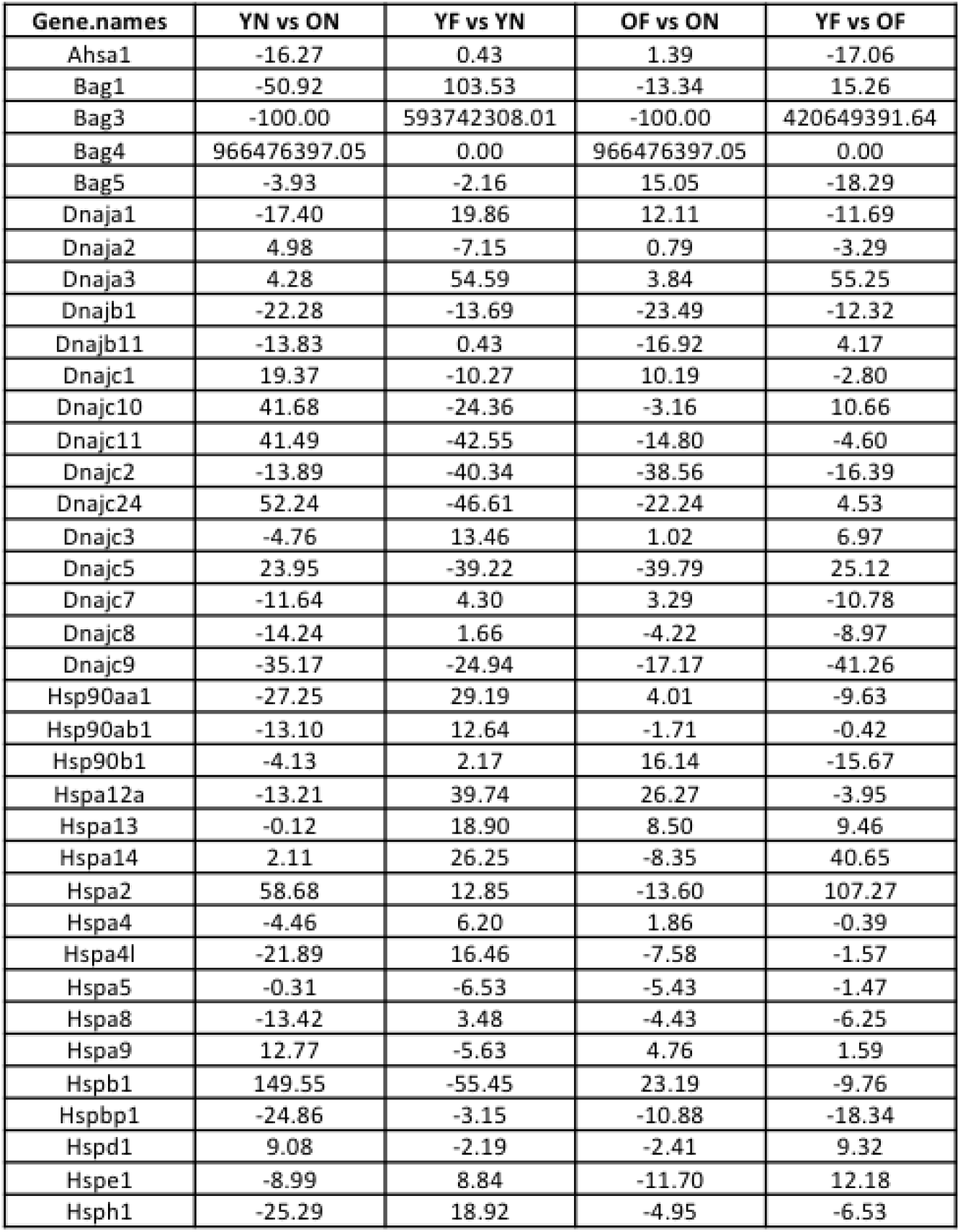
Table depicting the percent change in the expression of chaperome PN in alveolar macrophages (AM) across the indicated transitions.

**Table 2.** Table depicting the percent change in the expression of chaperome PN in alveolar macrophages (AM) across the indicated transitions.

| Gene.names | YN vs ON | YF vs YN | OF vs ON | YF vs OF |
| --- | --- | --- | --- | --- |
| Ahsa1 | -0.95 | 5.49 | 38.16 | -24.37 |
| Bag1 | -19.18 | 6.65 | -25.61 | 15.88 |
| Bag3 | -15.12 | -0.40 | -4.10 | -11.85 |
| Bag5 | -100.00 | 38.23 | 0.00 | -100.00 |
| Dnaja1 | -5.03 | -4.63 | -16.35 | 8.27 |
| Dnaja2 | -5.92 | -7.03 | -11.63 | -1.02 |
| Dnaja3 | 1.03 | 27.37 | 25.46 | 2.57 |
| Dnaja4 | -0.24 | -5.00 | 7.69 | -11.99 |
| Dnajb1 | -13.23 | -1.54 | -9.93 | -5.14 |
| Dnajb11 | -0.92 | -8.25 | -15.64 | 7.75 |
| Dnajb2 | 56.43 | -18.70 | 27.85 | -0.53 |
| Dnajb4 | 23.92 | -9.19 | 5.82 | 6.34 |
| Dnajc10 | 12.68 | -1.92 | -9.31 | 21.87 |
| Dnajc11 | -3.37 | 25.98 | 25.17 | -2.76 |
| Dnajc3 | -7.24 | -6.25 | -12.00 | -1.18 |
| Dnajc5 | 13.48 | -14.52 | 15.18 | -15.78 |
| Dnajc7 | -2.13 | -27.01 | -30.30 | 2.49 |
| Dnajc8 | -11.38 | 1.44 | 3.52 | -13.16 |
| Dnajc9 | -8.30 | -24.10 | -15.93 | -17.21 |
| Hsp90aa1 | -21.90 | 1.15 | -15.51 | -6.50 |
| Hsp90ab1 | -9.45 | -6.91 | -16.92 | 1.46 |
| Hsp90b1 | -7.22 | -0.96 | -8.83 | 0.80 |
| Hspa13 | 1.90 | -4.93 | -13.69 | 12.24 |
| Hspa1a | -38.45 | 49.34 | -19.15 | 13.69 |
| Hspa4 | 1.47 | -5.02 | -0.01 | -3.62 |
| Hspa4l | -25.90 | 19.14 | -5.92 | -6.16 |
| Hspa5 | -4.24 | -0.79 | -6.42 | 1.52 |
| Hspa8 | -9.58 | -6.14 | -14.95 | -0.21 |
| Hspa9 | -6.07 | 4.69 | 4.71 | -6.09 |
| Hspb1 | -56.67 | 102.53 | 18.23 | -25.78 |
| Hspd1 | -16.10 | 19.52 | 1.56 | -1.27 |
| Hspe1 | -10.53 | 85.82 | 19.45 | 39.18 |
| Hsph1 | -38.59 | 32.38 | -12.49 | -7.11 |

