## Supplemental Figures for "Proteostasis and Energetics as Proteome Hallmarks of Aging and Influenza Challenge in Pulmonary Disease"

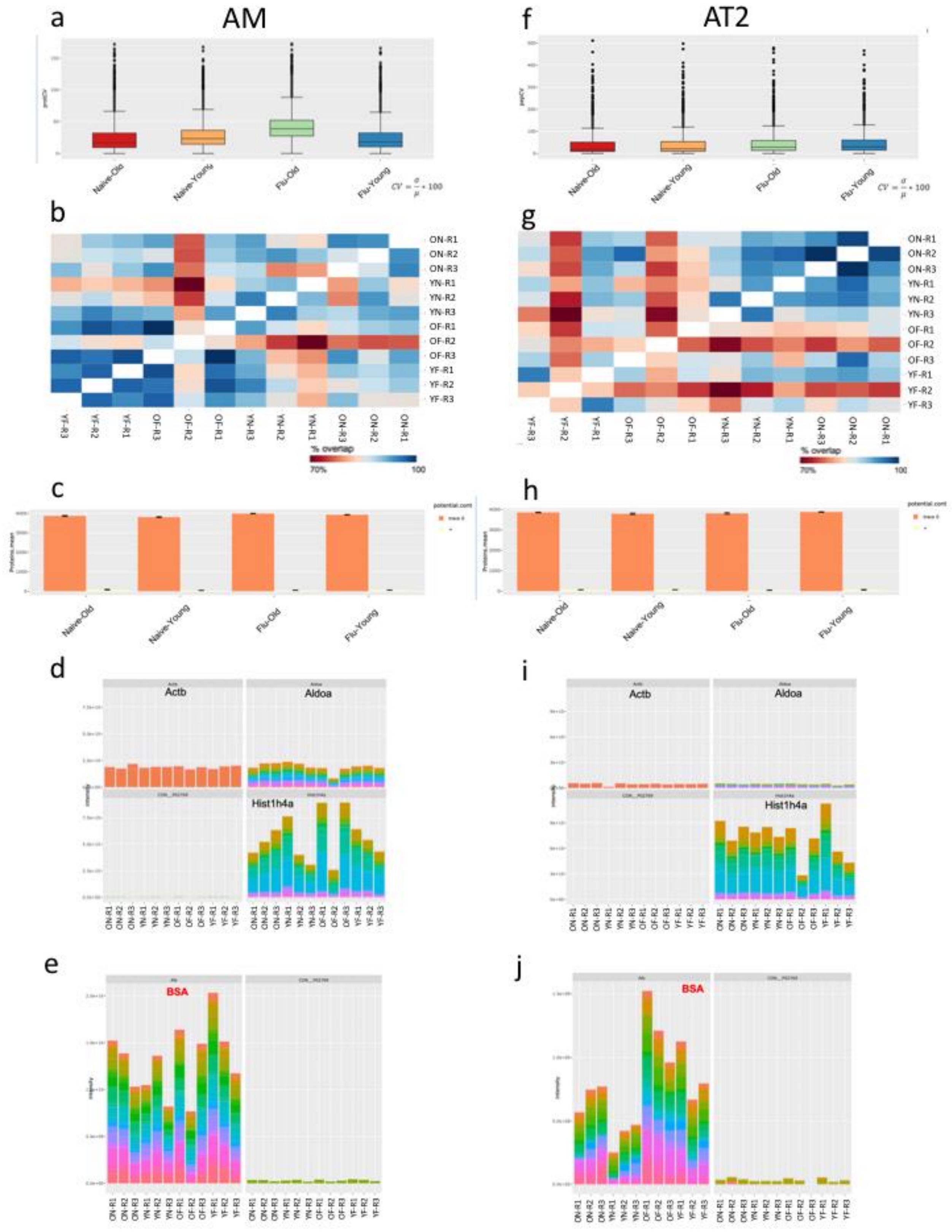
a
AM
Naive-Old
Naive-Young
Flu-Old
Flu-Young
$CV = \frac{\sigma}{\mu} * 100$
b
ON-R1
ON-R2
ON-R3
YN-R1
YN-R2
YN-R3
OF-R1
OF-R2
OF-R3
YF-R1
YF-R2
YF-R3
% overlap
70%
100
c
Naive-Old
Naive-Young
Flu-Old
Flu-Young
d
Actb
Aldoa
Hist1h4a
e
BSA
f
AT2
Naive-Old
Naive-Young
Flu-Old
Flu-Young
$CV = \frac{\sigma}{\mu} * 100$
g
% overlap
70%
100
h
Naive-Old
Naive-Young
Flu-Old
Flu-Young
i
Actb
Aldoa
Hist1h4a
j
BSA

**Figure S1. Quality control analysis reveals that the mass spectrometric data is high quality.** The coefficient of variation (CV) for protein intensities ($CV=(\sigma/\mu)*100$) is shown (**a & f**, AM and AT2 cells, respectively). With a median CV around 30%, and upper extreme well below 100%, the samples show a very good quality for each FACS sorted cell set. For comparison, similar CV distributions are usually obtained from samples analyzed using abundant cell culture samples. Pairwise overlap between replicates in terms of protein IDs (**b & g**) shows substantial overlap (80-100% range), with roughly four clusters corresponding to condition-age pairs. The barplots (**c & h**) show the total number of proteins identified, as averages and standard deviation (error bar) across replicates. A coverage of around 4000 is observed for all conditions, which is considered good for whole cell proteomics current standards. Moreover, such coverage is uniform and statistically significant across all conditions tested (**c & h**). Importantly, the number of potential contaminants (**c & h, yellow bars on the right**) is small related to recovered hits. In figs. **d & i**, intensities across conditions for abundant control proteins (Actb, Aldoa, Hist1h4a) serving as internal controls are consistent in total read. For both cell lines, the contribution of different replicates is comparable and proportional as expected. Representative contaminant protein intensities (labeled with CON_, lower left quadrant) are low. Finally, BSA contamination from contributed by FACS sorting is comparable across all samples, with the global effect of pushing proteins out of the “proteomics detection box” uniformly (**e-j**). The global effect of this residual contamination is estimated to push out ~300-400 low-abundance proteins outside detection.

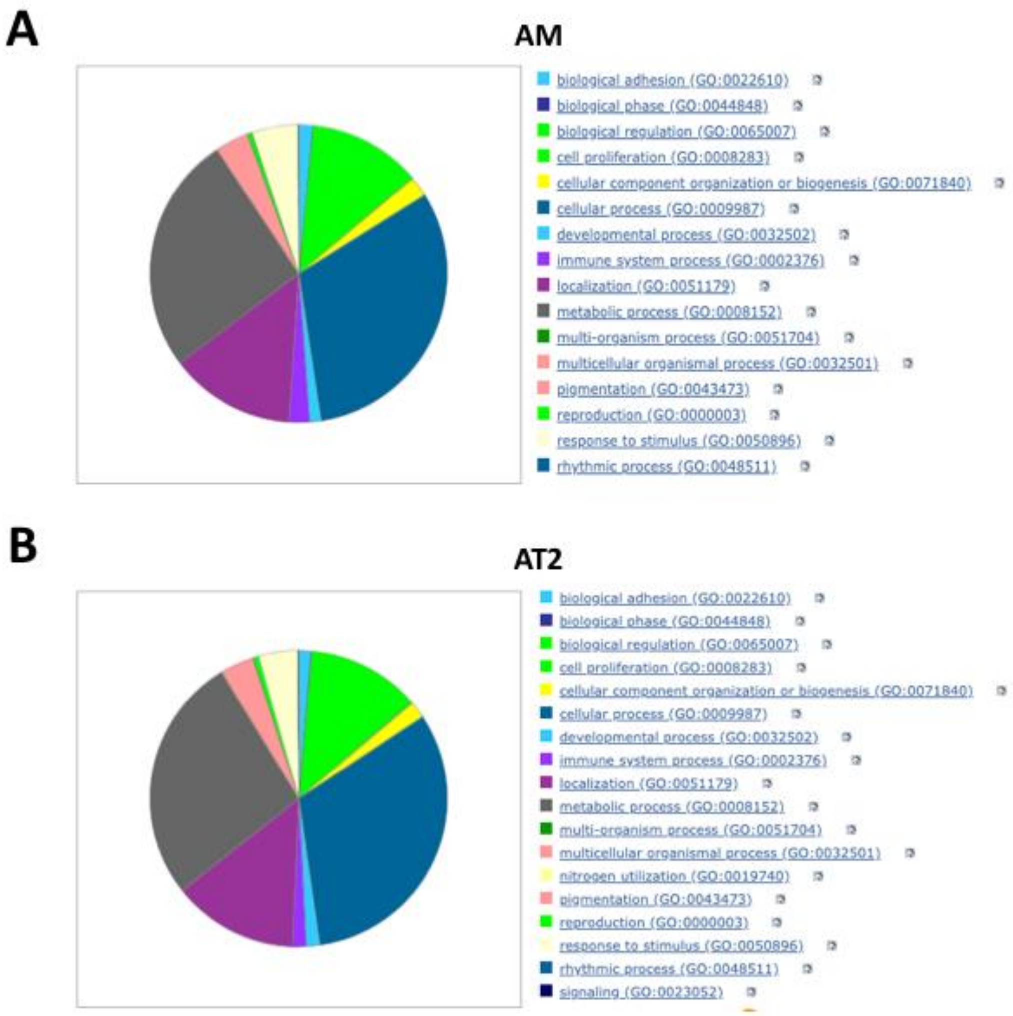


**Figure S2. GO Slim analysis of biological processes for the alveolar macrophage (AM) and alveolar type 2 (AT2) cell proteomes. A.** Pie chart highlighting the distribution of proteins recovered in AM lysates among the GO Slim Biological Processes. **B.** Pie chart highlighting the distribution of proteins recovered in AT2 lysates among the GO Slim Biological Processes.

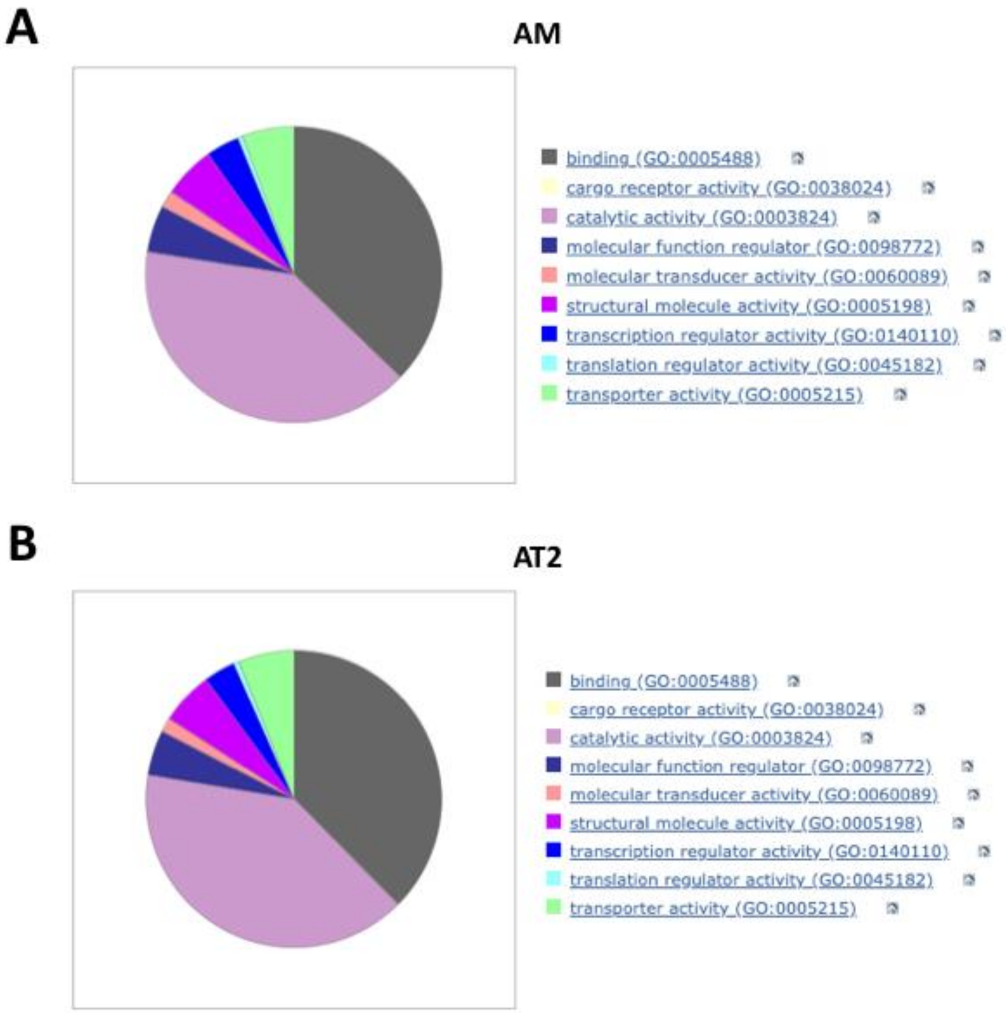


**Figure S3 GO Slim analysis of molecular functions for the alveolar macrophage (AM) and alveolar type 2 (AT2) cell proteomes. A.** Pie chart highlighting the distribution of proteins recovered in AM lysates among the GO Slim molecular functions. **B.** Pie chart highlighting the distribution of proteins recovered in AT2 lysates among the GO Slim molecular functions.

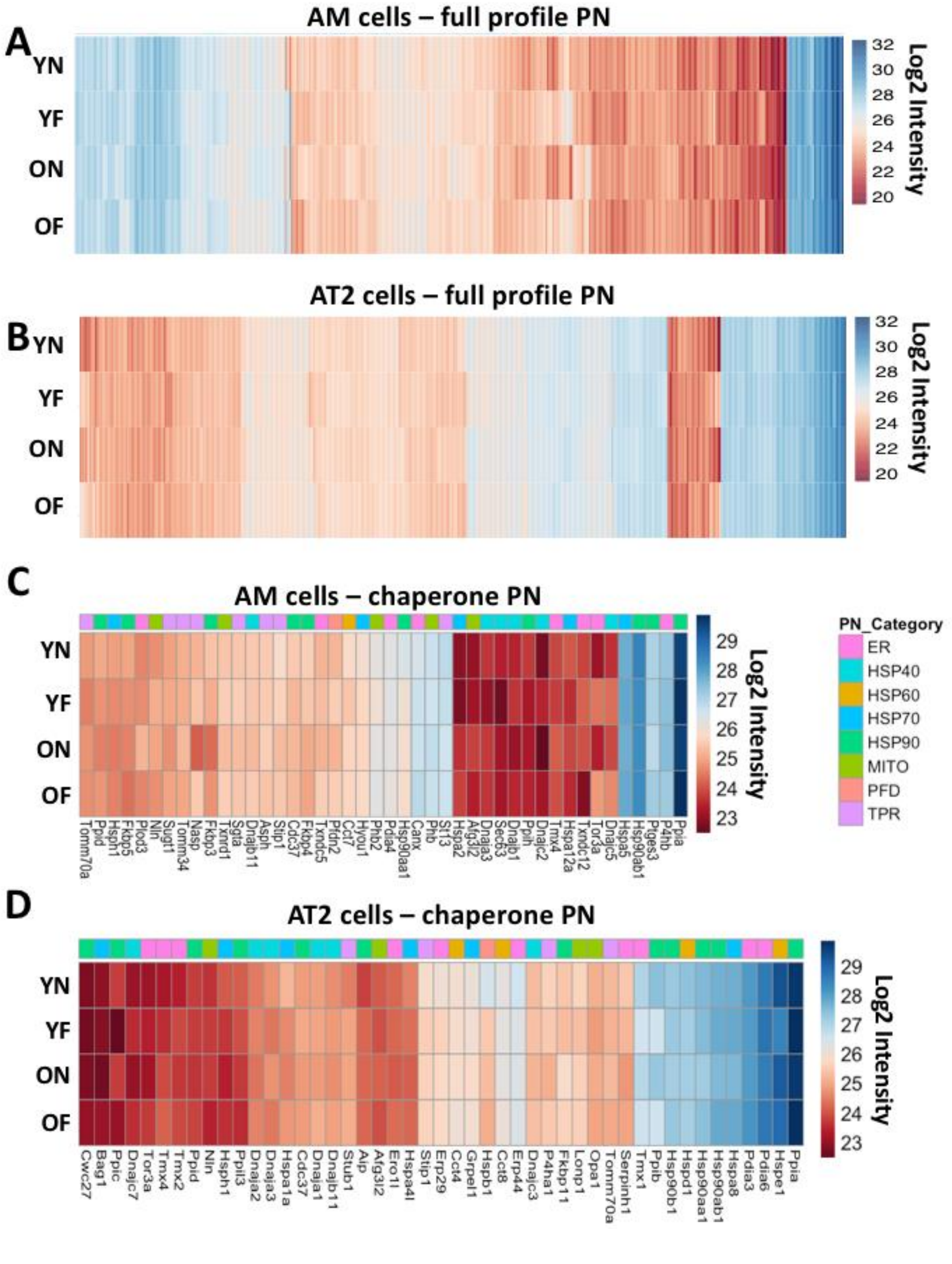
A
AM cells – full profile PN
YN
YF
ON
OF
32
30
28
26
24
22
20
Log2 Intensity
B
AT2 cells – full profile PN
YN
YF
ON
OF
32
30
28
26
24
22
20
Log2 Intensity
C
AM cells – chaperone PN
YN
YF
ON
OF
29
28
27
26
25
24
23
Log2 Intensity
Tomm70a Ppid Hsph1 Fkbp5 Plod3 Nln Sugt1 Tomm34 Nasp Fkbp3 Txnrd1 Sgta Dnajb11 Asph Stip1 Cdc37 Fkbp4 Txndc5 Pfdn2 Cct7 Hyou1 Phb2 Pdia4 Hsp90aa1 Canx Phb St13 Hspa2 Afg3l2 Dnaja3 Sec63 Dnajb1 Ppih Dnajc2 Tmx4 Hspa12a Txndc12 Tor3a Dnajc5 Hspa5 Hsp90ab1 Ptges3 P4hb Ppia
PN_Category
ER
HSP40
HSP60
HSP70
HSP90
MITO
PFD
TPR
D
AT2 cells – chaperone PN
YN
YF
ON
OF
29
28
27
26
25
24
23
Log2 Intensity
Cwc27 Bag1 Ppic Dnajc7 Tor3a Tmx4 Tmx2 Ppid Nln Hsph1 Ppil3 Dnaja2 Dnaja3 Hspa1a Cdc37 Dnaja1 Dnajb11 Stub1 Aip Afg3l2 Ero1l Hspa4l Stip1 Erp29 Cct4 Grpel1 Hspb1 Cct8 Erp44 Dnajc3 P4ha1 Fkbp11 Lonp1 Opa1 Tomm70a Serpinh1 Tmx1 Ppib Hsp90b1 Hspd1 Hsp90aa1 Hsp90ab1 Hspa8 Pdia3 Pdia6 Hspe1 Ppia

**Figure S4. Proteostasis components exhibit aged-specific and stress-specific expression patterns in alveolar cells. A.** Heat map depicting the log2 intensity of the expression of all identified full profile proteostasis network (PN) components in naïve (N) and influenza treated (F) young (Y) and old (O) alveolar macrophages (AM). **B.** Heat map depicting the log2 intensity of the expression of all identified full profile proteostasis network (PN) components in naïve (N) and influenza treated (F) young (Y) and old (O) alveolar type II cells (AT2). **C.** Heat map depicting the log2 intensity of the expression of all identified chaperone proteostasis network (PN) components in naïve (N) and influenza treated (F) young (Y) and old (O) alveolar macrophages (AM). **D.** Heat map depicting the log2 intensity of the expression of all identified chaperone proteostasis network (PN) components in naïve (N) and influenza treated (F) young (Y) and old (O) alveolar type II cells (AT2).

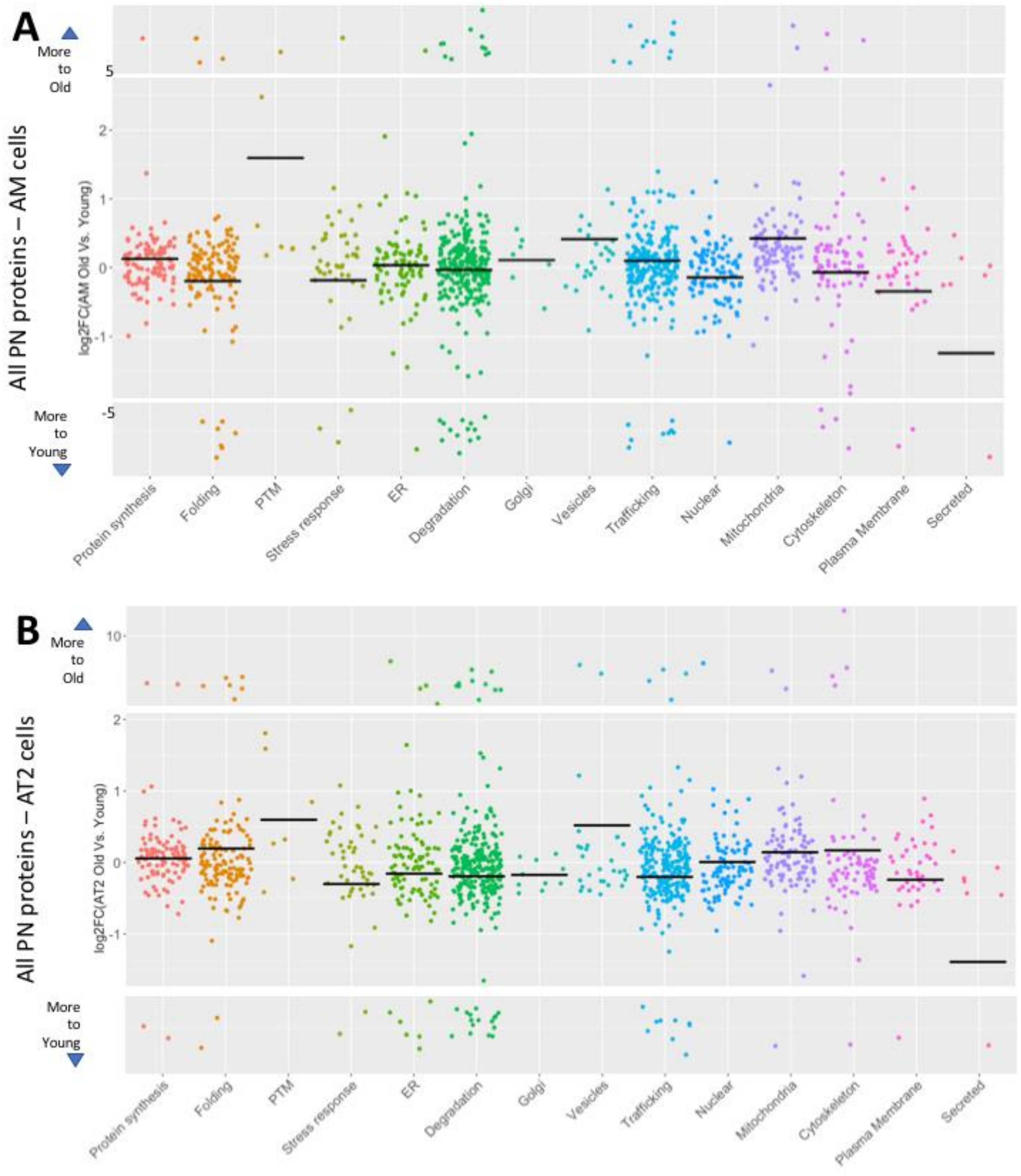

A
More to Old
More to Young
All PN proteins – AM cells
log2FC(AM Old Vs. Young)
5
2
1
0
-1
-5
Protein synthesis
Folding
PTM
Stress response
ER
Degradation
Golgi
Vesicles
Trafficking
Nuclear
Mitochondria
Cytoskeleton
Plasma Membrane
Secreted
B
More to Old
More to Young
All PN proteins – AT2 cells
log2FC(AT2 Old Vs. Young)
10
2
1
0
-1
Protein synthesis
Folding
PTM
Stress response
ER
Degradation
Golgi
Vesicles
Trafficking
Nuclear
Mitochondria
Cytoskeleton
Plasma Membrane
Secreted

**Figure S5. Proteostasis network (PN) components exhibit similar expression profiles between alveolar macrophages (AM) and type II (AT2) cells. A.** Scatter plot showing the log2 fold change (FC) of the average expression (n = 3) for each PN proteins identified in O versus Y AM cells (Log2 FC (O/Y)). **B.** Scatter plot showing the log2 fold change (FC) of the average expression (n = 3) for each PN proteins identified in O versus Y AT2 cells (Log2 FC (O/Y)). For both panels, the proteins are separated and colored according to their PN categorization displayed along the x-axis. Proteins with a log2 FC exceeding ± 2 were assigned a pseudo values of ± 10 and are displayed in the boxes at the top or bottom of the scatter plot. The bars represent the average of the expression for all proteins in the indicated PN category.

AM naive

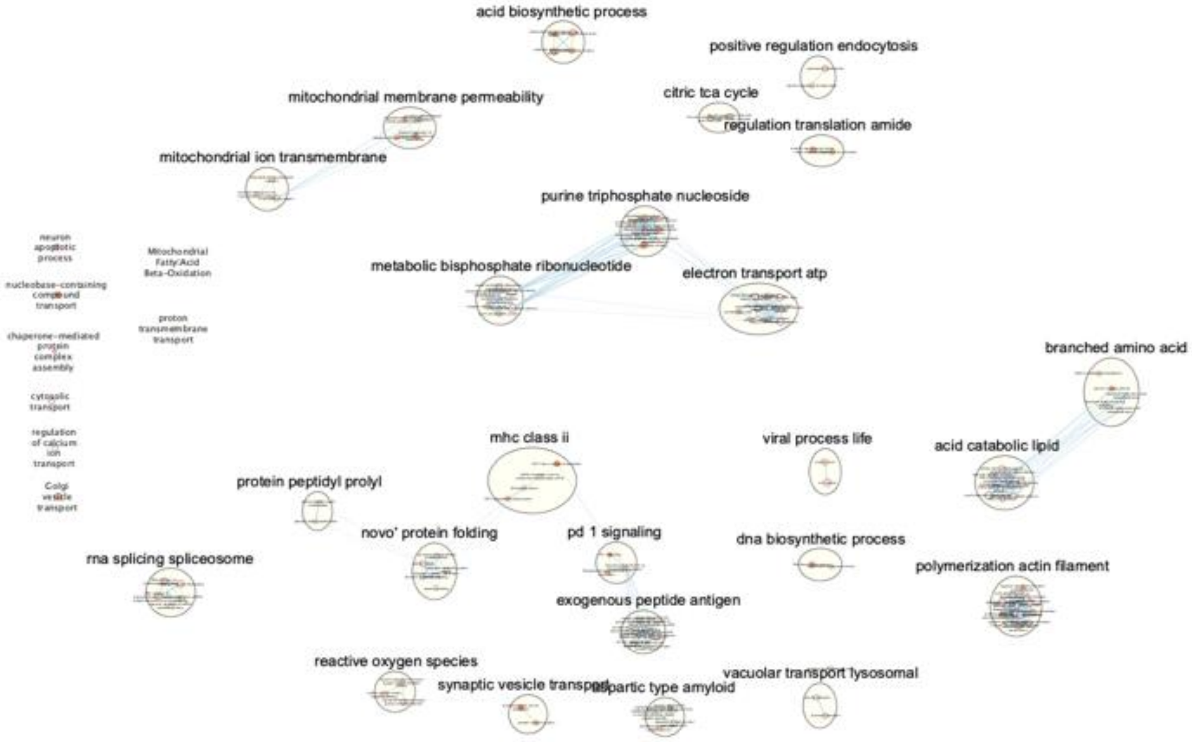


**Figure S6**

AM flu

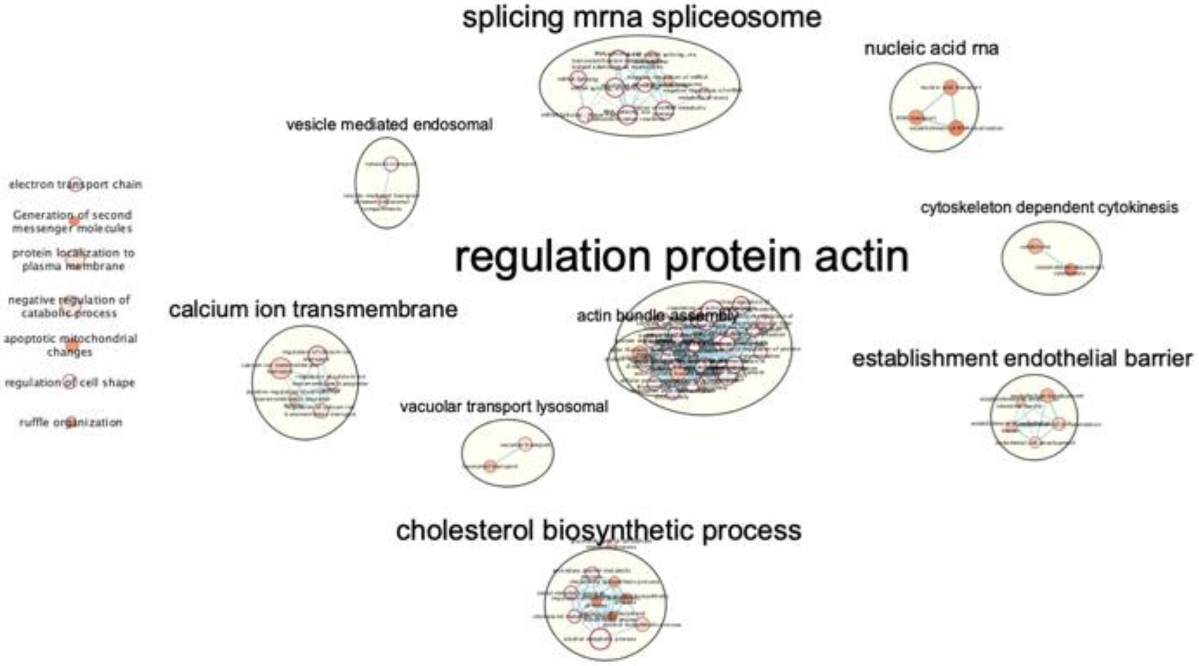


| Cluster | Nodes |
|---|---|
| regulation protein actin | 30 |
| splicing mrna spliceosome | 10 |
| cholesterol biosynthetic process | 9 |
| establishment endothelial barrier | 5 |
| calcium ion transmembrane | 5 |
| nucleic acid rna | 3 |
| vesicle mediated endosomal | 2 |
| vacuolar transport lysosomal | 2 |
| cytoskeleton dependent cytokinesis | 2 |
| actin bundle assembly | 2 |

**Figure S7**

AT2 naive

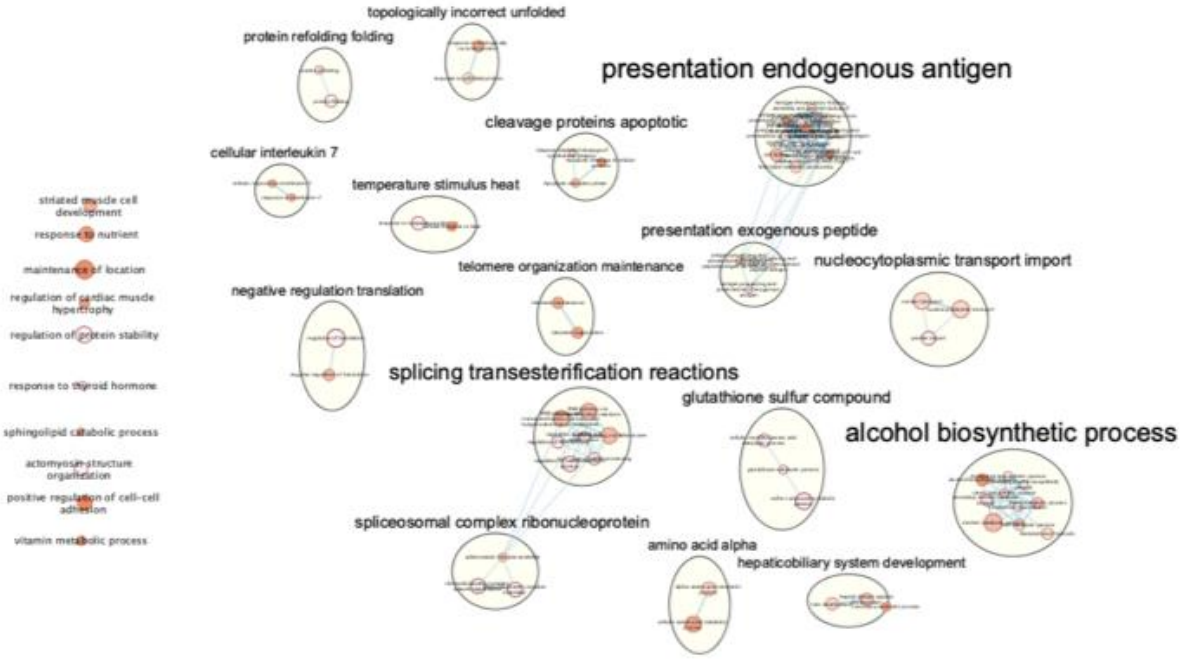


| Cluster | Nodes |
|---|---|
| presentation endogenous antigen | 13 |
| alcohol biosynthetic process | 10 |
| splicing transesterification reactions | 7 |
| spliceosomal complex ribonucleoprotein | 3 |
| presentation exogenous peptide | 3 |
| nucleocytoplasmic transport import | 3 |
| glutathione sulfur compound | 3 |
| cleavage proteins apoptotic | 3 |
| topologically incorrect unfolded | 2 |
| temperature stimulus heat | 2 |
| telomere organization maintenance | 2 |
| protein refolding folding | 2 |
| negative regulation translation | 2 |
| hepaticobiliary system development | 2 |
| cellular interleukin 7 | 2 |
| amino acid alpha | 2 |

**Figure S8**

AT2 flu

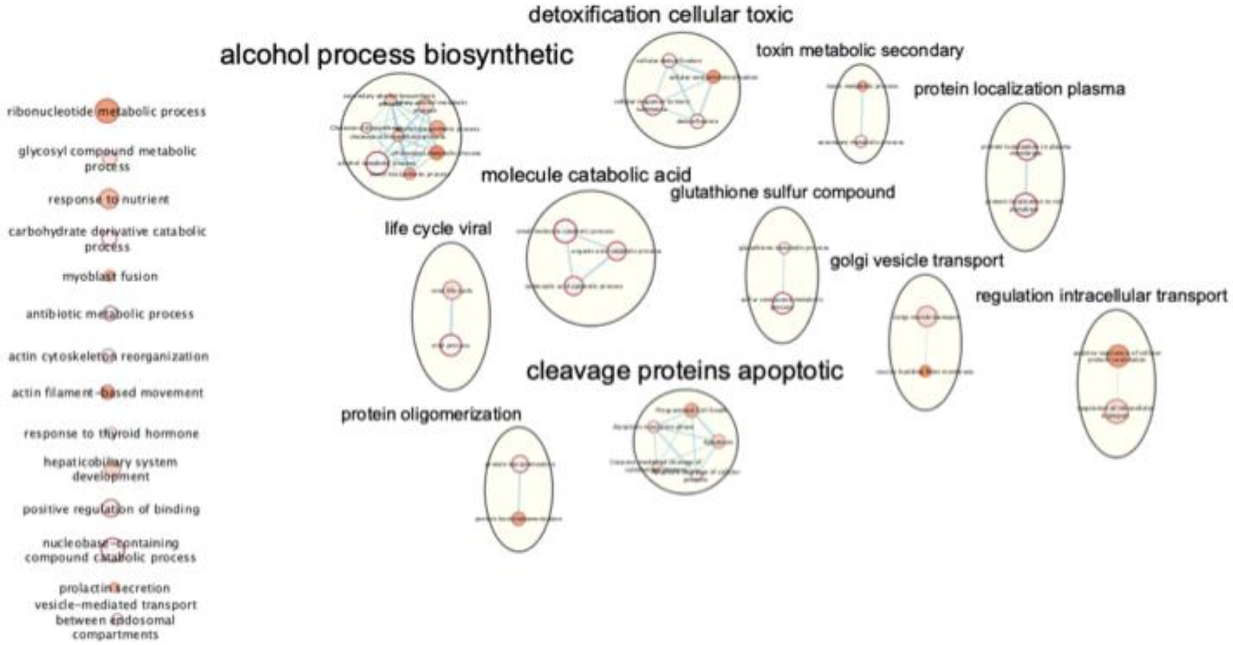


| Cluster | Nodes |
|---|---|
| alcohol process biosynthetic | 8 |
| cleavage proteins apoptotic | 5 |
| detoxification cellular toxic | 4 |
| molecule catabolic acid | 3 |
| toxin metabolic secondary | 2 |
| regulation intracellular transport | 2 |
| protein oligomerization | 2 |
| protein localization plasma | 2 |
| life cycle viral | 2 |
| golgi vesicle transport | 2 |
| glutathione sulfur compound | 2 |

**Figure S9**

**Figure S6-9**. **Enrichment Maps for GO::Biological Process and Reactome pathways, for AM YN-ON, AM YF-OF, AT2 YN-ON, AT2 YF-OF.** Each node (circle) represents a distinct pathway, and edges represent the number of proteins overlapping between two pathways, determined using a similarity coefficient. Individual node size is proportional to protein set size, and edge thickness is proportional to protein overlap size. Node color follows Q value (darker color->lower Q value). Enrichment maps were created with parameters FDR Q value <0.01 and combined similarity coefficient >0.375 with combined constant =0.5. In order to define major biological themes across the maps, clusters were automatically defined and summarized using the AutoAnnotate Cytoscape application. AutoAnnotate first clusters the network using the clusterMaker2 application and then summarizes each cluster on the basis of word frequency within the pathway names via a WordCloud application – here for each cluster showing the top three words corresponding to the most frequent node labels in the cluster.

| AM - GoSlim Biological Process | | | |
|---|---|---|---|
| Term | # proteins | % total | % Term total |
| cellular process | 1446 | 34.2% | 31.5% |
| metabolic process | 1194 | 28.3% | 26.0% |
| localization | 629 | 14.9% | 13.7% |
| biological regulation | 562 | 13.3% | 12.2% |
| response to stimulus | 230 | 5.4% | 5.0% |
| multicellular organismal process | 167 | 4.0% | 3.6% |
| immune system process | 102 | 2.4% | 2.2% |
| cellular component organization or biogenesis | 100 | 2.4% | 2.2% |
| biological adhesion | 70 | 1.7% | 1.5% |
| developmental process | 59 | 1.4% | 1.3% |
| reproduction | 26 | 0.6% | 0.6% |
| cell proliferation | 5 | 0.1% | 0.1% |
| biological phase | 3 | 0.1% | 0.1% |
| rhythmic process | 2 | 0.0% | 0.0% |
| multi-organism process | 1 | 0.0% | 0.0% |
| pigmentation | 1 | 0.0% | 0.0% |

| AT2 - GoSlim Biological Process | | | |
|---|---|---|---|
| Term | # proteins | % total | % Term total |
| cellular process | 1393 | 34.3% | 32.1% |
| metabolic process | 1170 | 28.8% | 26.9% |
| localization | 596 | 14.7% | 13.7% |
| biological regulation | 535 | 13.2% | 12.3% |
| response to stimulus | 188 | 4.6% | 4.3% |
| multicellular organismal process | 159 | 3.9% | 3.7% |
| cellular component organization or biogenesis | 80 | 2.0% | 1.8% |
| developmental process | 64 | 1.6% | 1.5% |
| immune system process | 63 | 1.6% | 1.4% |
| biological adhesion | 59 | 1.5% | 1.4% |
| reproduction | 25 | 0.6% | 0.6% |
| cell proliferation | 4 | 0.1% | 0.1% |
| biological phase | 3 | 0.1% | 0.1% |
| multi-organism process | 2 | 0.0% | 0.0% |
| pigmentation | 2 | 0.0% | 0.0% |
| nitrogen utilization | 1 | 0.0% | 0.0% |
| signaling | 1 | 0.0% | 0.0% |
| rhythmic process | 1 | 0.0% | 0.0% |

**Table S1.** Table depicting the GO Slim biological processes impacted by the identified proteins in alveolar macrophages (AM) (top) and alveolar type II cells (AT2) (bottom).

| AM - GoSlim Molecular Function | | | |
|---|---|---|---|
| Term | # proteins | % total | % Term total |
| catalytic activity | 1361 | 32.2% | 40.3% |
| binding | 1257 | 29.8% | 37.2% |
| structural molecule activity | 192 | 4.5% | 5.7% |
| transporter activity | 191 | 4.5% | 5.7% |
| molecular function regulator | 173 | 4.1% | 5.1% |
| transcription regulator activity | 122 | 2.9% | 3.6% |
| molecular transducer activity | 64 | 1.5% | 1.9% |
| translation regulator activity | 18 | 0.4% | 0.5% |
| cargo receptor activity | 2 | 0.0% | 0.1% |

| AT2 - GoSlim Molecular Function | | | |
|---|---|---|---|
| Term | # proteins | % total | % Term total |
| catalytic activity | 1286 | 31.7% | 40.0% |
| binding | 1205 | 29.7% | 37.5% |
| transporter activity | 195 | 4.8% | 6.1% |
| structural molecule activity | 187 | 4.6% | 5.8% |
| molecular function regulator | 159 | 3.9% | 4.9% |
| transcription regulator activity | 111 | 2.7% | 3.5% |
| molecular transducer activity | 50 | 1.2% | 1.6% |
| translation regulator activity | 18 | 0.4% | 0.6% |
| cargo receptor activity | 2 | 0.0% | 0.1% |

**Table S2.** Table depicting the GO Slim molecular functions impacted by the identified proteins in alveolar macrophages (AM) (top) and alveolar type II cells (AT2) (bottom).
